# A new therapeutic approach for Parkinson’s disease: dual targeting of α-Synuclein aggregation and microglial function by the novel immunomodulator 3-Monothiopomalidomide

**DOI:** 10.64898/2026.03.26.714051

**Authors:** Maria Francesca Palmas, Khosro Aminzadeh, Matteo Runfola, Pathik Parekh, Clara Porcedda, David Tweedie, Luca Casula, Maria Cristina Cardia, Jacopo Marongiu, Michela Etzi, Francesco Lai, Marcello Serra, Augusta Pisanu, Valeria Sogos, Alfonso De Simone, Dong Seok Kim, Nigel H. Greig, Anna R. Carta

## Abstract

**Background:** α-Synuclein (α-Syn) plays a central role in Parkinson’s disease (PD). Under pathological conditions, α-Syn aggregates into toxic oligomers and fibrils that act as damage-associated molecular patterns (DAMPs), stimulating microglial reactivity. This α-Syn–microglia axis creates a self-perpetuating cycle of neuroinflammation and neurodegeneration, accelerating dopaminergic neuron loss in the substantia nigra pars compacta (SNpc) and contributing to motor deficits. Moreover, α-Syn pathology spreads through the brain, disrupting synaptic plasticity in cognitive regions like the cortex and hippocampus, leading to early cognitive decline. Thus, targeting α-Syn aggregation and its inflammatory consequences presents a promising dual-hit therapeutic strategy for PD.

**Methods:** This study investigates the therapeutic potential of 3-monothiopomalidomide (3MP), a novel thalidomide derivative designed to reduce neuroinflammation with a potentially better safety profile than Pomalidomide (POM). The neuroprotective and anti-inflammatory effects of 3MP were evaluated in rat primary mesencephalic mixed neuron-microglia cultures exposed to human α-Syn oligomers (H-αSynOs). Anti-aggregation activity was assessed via Thioflavin T (ThT) assays and Thioflavin S (ThS) staining in SH-SY5Y cells. Finally, the anti-aggregation, anti-inflammatory, and neuroprotective effects of 3MP were evaluated *in vivo* in a rat model of PD induced by intracerebral infusion of H-αSynOs.

**Results:** In primary cell cultures, 3MP dose-dependently reduced α-Syn-induced neuronal death and microglial inflammatory responses. It also significantly inhibited α-Syn aggregation *in vitro* in the ThT assay and in SH-SY5Y cells exposed to α-Syn protofibrils, outperforming POM. When chronically administered *in vivo*, 3MP preserved dopaminergic neurons within the SNpc and yielded functional benefits on motor and cognitive readouts. Notably, 3MP markedly attenuated α-Syn aggregates induced by the H-αSynOs infusion in the SNpc more efficiently than POM, as shown by reduced intraneuronal staining for pSer129-α-Syn+ and reduced pSer129-αSyn in both cytoplasmic and phagolysosomal compartments of microglia. In addition, mesencephalic and cortical inflammatory microgliosis that followed to intranigral H-αSynOs-infusion, were significantly dampened by 3MP.

**Conclusions:** Overall, 3MP emerges as a dual-action drug candidate capable of modulating neuroinflammation and α-Syn aggregation and thereby disrupting the α-Syn-driven inflammatory cycle. Its neuroprotective effects and favourable safety profile support its potential as a disease-modifying therapy for PD, with promising implications for clinical translation.

## Background

The recognition of α-synuclein’s (α-Syn) central role in Parkinson’s disease (PD) pathophysiology [1] has redefined PD as a misfolding proteinopathy, establishing a framework to classify neurodegenerative disorders by their dominant pathological protein aggregates. Under physiological conditions, α-Syn predominantly exists as disordered monomers or α-helical multimers [2,3]. However, under pathological conditions, α-Syn converts into various toxic species, including oligomers, protofibrils, and fibrils. Among these, oligomers enriched in β-sheet structures are considered the most neurotoxic due to their ability to disrupt neuronal membranes, impair proteostasis [4] and pathologically interact with glial cells to trigger neuroinflammatory responses [5,6]. Importantly, α-Syn oligomers serve as seeds for aggregate propagation through prion-like mechanisms [7,8]. Therefore, addressing the α-Syn aggregation process has become a pivotal, yet unmet objective in drug design for PD [9–11].

Neuroinflammation is a complex process within the central nervous system (CNS), primarily mediated by microglia, that serves both adaptive and maladaptive roles. While neuroinflammation protects against injuries or infections, a chronic and/or dysregulated microglial response leads to excessive release of proinflammatory mediators, which contribute to neuronal dysfunction and damage [8]. This persistent neuroinflammatory response, initially beneficial, becomes maladaptive over time and is a hallmark of many neurodegenerative diseases, including PD [12–15]. In PD, the chronic neuroinflammatory response is prominent in the substantia nigra pars compacta (SNpc), a region where dopaminergic neurons display heightened vulnerability to inflammatory insults [16–19]. This localized inflammation significantly contributes to the degeneration of these neurons, thereby playing a key role in the emergence of the typical motor symptoms of the disease.

In addition to its well-characterized motor manifestations, PD is frequently accompanied by cognitive disturbances, which can appear early in the disease course as mild cognitive impairment (MCI) and may later progress to Parkinson’s disease dementia (PDD) [20,21]. While cognitive decline in advanced stages of PD has been associated with widespread Lewy body (LB)-pathology, particularly in cortical areas [22], it is noteworthy that MCI is also common in early or newly diagnosed cases, often preceding the detectable accumulation of neocortical LB deposits [20,23,24]. This temporal discrepancy raises important questions about the mechanisms driving early cognitive changes and suggests that pathological alterations might begin before α-Syn aggregation becomes histologically apparent. Increasing evidence points to neuroinflammation as a potential contributor, particularly within brain regions involved in cognition, such as the hippocampus and anterior cingulate cortex (ACC) [25–27]. In these areas, reactive microglia release proinflammatory cytokines that interfere with synaptic function and plasticity, ultimately disrupting neuronal communication and facilitating cognitive decline.

Established that the bidirectional interplay between α-Syn pathology and microglial reactivity drives a self-perpetuating cycle of neurodegeneration, it becomes pivotal to underscore immunomodulation as a therapeutic priority. In this context, immunomodulatory imide drugs (IMiDs) hold significant promise due to their unique ability to reduce neuroinflammation and rescale immune responses in cellular and animal models of neurological disorders [28–31]. Thalidomide-like IMiD compounds have shown considerable potential in decreasing inflammatory cytokine production, particularly tumour necrosis factor (TNF)-α, a master regulator of innate immune responses. By lowering TNF-α levels, these drugs modulate the inflammatory process, offering neuroprotective benefits in conditions such as PD, Alzheimer’s disease (AD), and other neurodegenerative disorders where chronic inflammation accelerates neuronal damage [32]. Notably, our previous research has demonstrated that the third-generation IMiD Pomalidomide (POM), not only reverses PD-associated motor deficits but also protects against nigral dopaminergic cell loss induced by intracerebral infusion of toxic α-Syn. Moreover, POM effectively modulates the inflammatory response both in the CNS and peripherally [33]. Current immunomodulatory therapies, while effective, are often limited by significant teratogenic risks. This underscores the growing need for new drugs that can retain therapeutic efficacy while minimizing adverse developmental effects. A recent study has highlighted 3-monothiopomalidomide (3MP), a novel thalidomide derivative, as a promising candidate in this regard. 3MP effectively suppresses TNF-α production and broader neuroinflammatory responses, as demonstrated in a mouse model of traumatic brain injury (TBI), where it reduced astroglial and microglial activation and improved behavioral outcomes. Additionally, 3MP binds to human cereblon (CRBN) without inducing the degradation of Sal-like protein 4 (Sall4), a pathway implicated in the teratogenicity of compounds such as POM [34]. Importantly, 3MP lacked toxicity in chicken embryos [34], a classical model for early teratogenicity assessment.

Building on these findings, in the present study we evaluated the therapeutic potential of 3MP, aiming to further characterize its efficacy in the context of α-Syn neuropathology, neuroinflammation and secondary damage associated with PD. The neuroprotective and anti-inflammatory potential of 3MP was initially screened *in vitro* using rat primary mesencephalic neurons and microglia exposed to α-Syn. The compound’s anti-aggregation activity was then assessed both in acellular systems, via the Thioflavin T (ThT) aggregation assay, and in cellular models using Thioflavin S (ThS) staining. Subsequently, the drug was evaluated for its anti-aggregant, anti-inflammatory and neuroprotective properties when chronically administered *in vivo* as a nanosuspension [35], in a rat model of α-synucleinopathy induced by the intracerebral infusion of exogenous human α-Syn oligomers (H-αSynOs), previously characterized and validated for PD-like neuropathology and behavioral phenotype [33,36]. Collectively, our results propose 3MP as a dual-action therapeutic candidate capable of preventing α-Syn aggregation and disrupting the α-Syn/neuroinflammation cycle by restoring microglia function via its potent immunomodulatory actions, to ultimately mitigate α-Syn neuropathology and providing first evidence of functional benefit in our *in vivo* disease model.

## Materials and methods

### Expression and purification of recombinant α-Syn

Recombinant human N-terminally acetylated α-Syn was expressed in BL21-Gold (DE3) *E. coli* competent cells (Agilent #230132) by co-transforming a plasmid encoding the α-Syn sequence with one expressing the NatB complex (naa20 and naa25 genes). In particular, the pT7–7 plasmid has been used in association with ampicillin resistance to express human α-Syn while the pACYCduet-naa20-naa25 plasmid in association with chloramphenicol resistance has been used to express NatB. This established protocol ensures 100% N-terminal acetylation of α-Syn [4]. Following co-transformation, cells were plated on LB agar with ampicillin (100 μg/mL) and chloramphenicol (12.5 μg/mL). A single colony was selected and cultured at +37 °C, 180 RPM. The co-transformed cells were induced at OD□□□ = 0.6 by the addition of 1 mM isopropyl β-d-1-thiogalactopyranoside (IPTG) at 37 °C for 4 h. Cells were centrifuged (6200□*g* for 50min) and resuspended in lysis buffer (10 mM Tris-HCl pH 8, 1 mM EDTA, EDTA-free complete protease inhibitor cocktail tablets obtained from Roche, Basel, Switzerland) and lysed by sonication. The cell lysate was centrifuged at 22,000g for 30 min to remove cell debris. In order to precipitate the heat-sensitive proteins, the supernatant was then heated for 20 min at 80 °C and centrifuged at 22,000g. Subsequently streptomycin sulphate was added to the supernatant to a final concentration of 10 mg.ml-1 to stimulate DNA precipitation. The mixture was stirred for 15 min at 4 °C followed by centrifugation at 22,000g. Then, ammonium sulphate was added to the supernatant to a concentration of 360 mg.ml-1 to precipitate the protein. The solution was stirred for 30 min at 4 °C and centrifuged again at 22,000g. The resulting pellet was resuspended in 25 mM Tris-HCl, pH 7.7 and dialyzed against the same buffer to remove salts. After filtration (0.22 μm), the sample was loaded onto a Q Sepharose anion exchange column and eluted with a 0–1 M NaCl gradient in 25 mM Tris-HCl (pH 7.7) buffer. Selected fractions were pooled together and further purified via size-exclusion chromatography (Superdex 75, 26/600) in PBS (pH 7.4). Protein fractions have been subjected to SDS-PAGE to prove purification efficiency and finally concentrated using 10 kDa MWCO centrifugal filters (Sartorius Stedim Biotech, Gottingen, Germany). The protein concentration was determined from the absorbance at 275 nm using an extinction coefficient of 5600 M^−1^ cm^−1^. The fraction has been aliquoted (1.0 mL, 280 μM) and finally flash-frozen in liquid nitrogen, stored at −20 °C until use.

### Drug synthesis and preparation

3MP was synthesized from POM by thionation with phosphorus pentasulfide through the intermediate 3,6’-dithiopomalidomide. Resulting 3MP was purified by prep-HPLC to yield an orange-red powder.

3MP was formulated as a nanosuspension by wet ball media milling following a reported procedure [35,37]. Briefly, the drug (1% (w/v)) was, at first, dispersed in a tween 80 aqueous solution (0.5% w/v) by vortexing, and then milled for 60 minutes in Eppendorf microtubes containing yttrium-stabilized zirconia-silica beads (Silibeads® Type ZY Sigmund Lindner, Germany) with a diameter of 0.1–0.2 mm using a beads-milling cell disruptor equipment (Disruptor Genie®, Scientific Industries, USA). The beads were removed by sieving, and the final formulation was analysed using a Zetasizer nano (Malvern Instruments, Worcestershire, UK) to verify the nanocrystal’s dimensional properties (average diameter and polydispersity index) and zeta potential.

### Cellular studies

Rat dopaminergic neurons and microglia were cultured as described by Zhang et al., 2005 [38] and Callizot et al, 2019 [39], with modifications. Briefly, the ventral segment of the mesencephalic flexure, a midbrain region abundant in dopaminergic neurons, was aseptically removed from euthanized rat fetuses (14-day gestation) [38,39] and were incubated for 20 min at 37 °C with a trypsin/EDTA solution (0.05 % trypsin and 0.02 % EDTA). Dulbecco’s modified Eagle’s medium (DMEM) containing DNAase I grade II (0.5 mg/mL) and 10 % of fetal calf serum (FCS) was added, and cells were mechanically dissociated by 3 passages through a 10 ml pipette. Following centrifugation (180 x g, 10 min, 4°C on a layer of BSA (3.5 %) in L15 medium), the cell pellet was suspended in culture medium (neurobasal medium with a 2 % solution of B27 supplement, 2 mmol/liter of L-glutamine, 2% PS solution, 10 ng/ml of brain-derived neurotrophic factor (BDNF), 1 ng/ml of glial cell line-derived neurotrophic factor (GDNF), 4% heat-inactivated FCS, 1g/L of glucose, 1 mM of sodium pyruvate, and 100 µM of non-essential amino acids). Cells were ultimately seeded (80,000 cells/well in 96 well-plates (poly-L-lysine coated)) and placed in an incubator (37 °C, 5% CO2, 95 % air). Half of the medium was replenished every other day.

At day 7, cells were incubated for 1 hr with culture medium freshly spiked with known concentrations of 3MP or vehicle and then challenged with α-Syn (72 hr, 250 nM α-syn (prepared as a 4 mM solution and pre-incubated 37o C for 72 hr in the dark with shaking to induce oligomerization)). Culture supernatant was removed and frozen for later analysis. Cells were washed (PBS) and then fixed (4% paraformaldehyde in PBS (pH 7.3, 21o C, 20 min)), washed (PBS) twice, and permeabilized. PBS containing 0.1 % of saponin and 1% of FCS was added (21o C, 15 min) to remove non-specific binding, and cells were then incubated with either (i) rabbit polyclonal anti tyrosine hydroxylase (TH) (1:2000 in PBS containing 1% FCS, 0.1% saponin, 21o C, 2 hr) to visualize dopaminergic neurons and neurites, or (ii) mouse monoclonal anti-IBA1 (1:500 in PBS containing 1% FCS and 0.1 % of saponin, 21o C, 2 hr) to visualize microglia. Cells were washed and incubated with Alexa Fluor® 488 goat anti-rabbit IgG (Jackson, 1:400) in PBS containing 1% FCS, 0.1% saponin (21o C, 1 hr.) Photomicrographs (20/well x 10 magnification) were acquired (ImageXpress, Molecular Devices, San Jose, CA) and analyzed by Custom Module Editor® (Molecular Devices). TNF-α protein levels were quantified by ELISA (rat TNF-α ELISA, Abcam, ab46070).

The evaluated concentration range of 3MP (3 to 60 µM) was selected from prior studies demonstrating the tolerability of 3MP and close analogues across immortal and primary cell cultures [31,32,34]. The selected concentration of α-Syn challenge (250 nM) was chosen from prior studies demonstrating its induction of a substantial and statistically significant proinflammatory response [38,39].

### *In vitro* aggregation experiments

To monitor α-Syn aggregation we used an established assay based on the fluorescence of ThT, which is induced when the dye binds to amyloid aggregates. α-Syn aggregation was induced in PBS buffer using 2.5 mM CaCl□ and a protein concentration of 100 μM. The solution also included 0.02% sodium azide (w/v) as bacteriostatic and 50 μM ThT. All experiments here performed also included 2% DMSO, to solubilize POM and 3MP when needed. The samples solutions were transferred into wells in triplicates (100 μL per well) using 96well plates (Grenier 655161), covered with aluminium foil to prevent solvent evaporation (ThermowellTM Sealing Tape - Corning). To monitor the aggregation kinetics via ThT fluorescence, we employed a FLUOStar Omega (BMG Labtech) plate reader with excitation and emission wavelengths set at 448 nm and 482 nm, respectively. 720 cycles, with cycle times of 600 seconds and 3 flashes per reading in each well, were employed for a total time of 120h. Incubations were performed at 37 C°, with orbital shaking at 200 RPM between the reading. The kinetic data obtained from the aggregation assays were normalized with respect to maximum ThT fluorescence intensities of the control protein-alone sample. The data have been analyzed and plotted with the Prism program.

Plateau and half-time (t½) values were determined for both treatments and for the control (protein alone). Plateau values were defined as the average of ThT fluorescence measured between 96 and 120 hours. The aggregation half-time (t½) was defined as the time needed for the ThT fluorescence intensity to reach 50% of its maximum (plateau) value.

### TEM imaging

TEM images were measured on the samples after 120h incubation. 5□μL of each sample were applied onto a carbon-coated 200 mesh copper. Following adsorption, the sample was washed twice with water and negatively stained with phosphotungstic acid (PTA) (2% w/v) for 2 min. TEM images were obtained using a Tecnai G2 S-TWIN microscope (Thermo Fisher Scientific) operating with a Eagle4K camera. Each condition was imaged at 100 nm and 200 nm magnifications.

### *In vitro* aggregation assay on SH-SY5Y cells

SH-SY5Y human neuroblastoma cells (obtained from ICLC Cell bank) were seeded on coverslips and grown in high glucose DMEM supplemented with 10% fetal bovine serum (FBS), 100 units/mL of penicillin and 100 µg/mL of streptomycin and maintained at 37 °C in a humid atmosphere with 5% CO2. Cells were pre-treated for 1 hour with 60 μM of POM or 3MP, followed by the addition of 1 μM of α-Syn pre-formed fibrils (PFFs; StressMarq Biosciences Inc., #SPR-317) to the culture medium. After four days of incubation, SH-SY5Y cells were fixed in 4% paraformaldehyde for 30 minutes, washed twice with PBS and permeabilised with PBS-Triton 0.1%. To detect α-Syn aggregates, coverslips with adherent cells were incubated for 15 minutes in a 0.05% ThS solution prepared in 50% ethanol/water, washed twice with 50% ethanol/water, once with 80% ethanol, briefly rinsed with water, and mounted on glass slides. Images were acquired using a spinning disk confocal microscope (Crisel Instruments, Rome, Italy) at a magnification of ×63.

### Animals, stereotaxic surgery, and pharmacological treatment

All experiments were performed in accordance with the ARRIVE guidelines and with the guidelines approved by the European Community (2010/63UE L 276 20/10/2010). Experimental procedures were approved by the Italian Ministry of Health (authorization n° 766/2020-PR). Every effort was made to minimize pain and discomfort and to reduce the number of animals used in the experiments.

Male Sprague Dawley rats (275–300 g, approximately 3 months old) were purchased from Envigo. All rats were housed in groups of four in polypropylene cages, with food and water available ad libitum, in rooms maintained at 21□°C under a 12□h light/dark cycle (lights on 7:00 A.M.). Prior to surgery, animals were deeply anesthetized with fentanyl (0.33 mg/kg, i.p.) and medetomidine hydrochloride (0.33 mg/kg, i.p.) and positioned in a stereotaxic apparatus (DKI-900LS, David Kopf Instruments, Tujunga, CA). Under aseptic conditions, a midline incision was made to expose the skull. Stereotaxic injections of 5 µL H-αSynOs per side were administered into the SNpc at a rate of 1 µL/min using a silica microinjector, as previously described [33,36] (coordinates relative to bregma: −5.4 mm anteroposterior; ±1.9 mm mediolateral; −7.2 mm dorsoventral, according to the atlas of Paxinos and Watson) [40]. Control animals received an equivalent volume of sterile phosphate-buffered saline (PBS, pH 7.4) at the same infusion rate. After inoculation, the cannula was kept in place for an additional 5 minutes and then gradually withdrawn to prevent backflow along the needle track.

To support dose-selection for the chronic treatment, a preliminary tolerability study was conducted in healthy age-matched rats. In this pilot experiment, animals received 3MP intraperitoneally for 2 months either at 5 mg/kg on alternate days (three times per week) or at 10 mg/kg once daily. Body weight was recorded longitudinally, and motor performance was assessed. Moreover, livers were collected from all rats to evaluate potential 3MP–related hepatic toxicity. For this purpose, a classical haematoxylin-eosin staining was applied, and sections were evaluated by a pathologist.

One-month post-surgery, animals were subjected to pharmacological treatments consisting of either the 3MP nanosuspension or a saline solution for 2 months and sacrificed 24 h after the last injection. Based on the pilot tolerability study, the experimental protocol included the evaluation of two distinct doses with separate administration regimens: 5 mg/kg (i.p.) administered on alternate days, three days per week, and 10 mg/kg (i.p.) administered daily. Notably, a nanosuspension specifically optimized for 3MP was used, which displayed a more favourable pharmacokinetic profile than the coarse suspension [35].

To compare 3MP with the third-generation IMiD POM in targeting α-Syn aggregation, a second group of animals was treated with the latter compound, which was also formulated as a nanosuspension following our previous protocol [37]. Specifically, POM was administered at a dose of 20 mg/kg, three times per week, as previously shown [33].

### Behavioral testing

Motor and non-motor behavioral tests were conducted over a 7-day period, starting 12 weeks after HαSynOs infusion. To minimize alterations in behavioral parameters due to the novel environment, rats were acclimated to the testing room for 30 minutes prior to each test. Behavioral assessments were performed between 9:00 a.m. and 3:00 p.m. All tests were conducted and analyzed by individuals blinded to the experimental conditions

#### Challenging beam walk test

Motor coordination and balance were evaluated by means of the challenging beam test following previously established methodologies [33,36], with adaptations from Drucker-Colín and García-Hernández [41], Fleming et al.[42], and Korecka et al. [43]. Briefly, the apparatus consisted of a 2-meter wooden beam set at a 15° incline, connecting a starting platform elevated 40 cm above the floor to the home cage. Beams of three different widths (15 mm, 10 mm, and 5 mm) were utilized. All rats underwent training to traverse the beams over a 3-day period before testing. On the test day, the sessions were videotaped for analysis.

During testing, each rat was placed at the lower end of the beam, and the number of stepping errors made while crossing toward the home cage was recorded. This procedure was repeated for all three-beam width. In cases where an animal failed to complete the task within the allocated time or fell off the beam, this was interpreted as an additional sensorimotor impairment, consistent with prior reports [33]. To quantify the degree of impairment, an incremental error score was assigned based on the following criteria: (A) a 0.25 increment if the animal completed 75% of the beam, (B) a 0.5 increment if 50% of the beam was traversed, and (C) a 0.75 increment if only 25% of the beam was completed.

#### Novel object recognition

The Novel Object Recognition (NOR) test was performed in a black box (60 × 60 cm) following a previously published protocol [27]. After an initial habituation session (10 minutes, T0), rats were individually reintroduced into the test box, where they were presented with two identical objects for 10 minutes before being returned to their home cage (familiarization phase, T1). Sixty minutes later, the rats were placed back in the test chamber for 3 minutes, where one familiar object and one novel object were presented (choice phase, T2). Training and testing sessions were recorded using a camera. Object recognition was quantified using the discrimination index (DI), calculated as follows: (Tn − Tf) / (Tn + Tf), where Tn represents the time spent exploring the novel object and Tf represents the time spent exploring the familiar object.

### Immunohistochemistry

After behavioral testing and 24 hours following the final injection, rats were deeply anesthetized and transcardially perfused with ice-cold 0.1 M PBS (pH 7.4) followed by 4% paraformaldehyde. Following perfusion, the brains were carefully removed from the skull, post-fixed overnight in 4% paraformaldehyde-PBS, and stored in 0.1% NaN3-PBS at 4°C. Serial coronal sections of the ACC and midbrain were cut using a vibratome (40 µm thickness) (Leica VT1000 S, Leica Biosystems, Germany). For TH neuronal counting, midbrain sections were pre-incubated with normal donkey serum (NDS) and then incubated with a rabbit polyclonal anti-TH primary antibody (1:1000, Millipore, Burlington, MA, USA). The reaction was amplified using a biotinylated secondary antibody and visualized using the avidin-biotin-peroxidase complex (ABC, Vector Laboratories, UK) method, with 3,3’-diaminobenzidine (DAB, Sigma-Aldrich, St. Louis, MO, USA) as the chromogen. Sections were then mounted on microscope slides, dehydrated, and coverslipped using Eukitt mounting medium (Fluka, Sigma-Aldrich, Italy), and allowed to dry overnight. Images were acquired with a Zeiss Axio Scan Z1 slide scanner (Zeiss, Germany) with a 20x magnification.

For immunofluorescence, midbrain and cortical sections were blocked with 1% BSA, 10% NDS and 0.1% Triton X-100 in PBS at 4□°C overnight. The day after, sections were immunoreacted with the following unconjugated primary antibodies for double immunolabeling: rabbit monoclonal anti-pSer129-αSyn (1:800, Abcam, Cambridge, UK); mouse monoclonal anti-TH (1:400, Millipore, Burlington, MA, USA); goat polyclonal anti Iba-1 (1:1000; Novus Biologicals, Littleton, CO, USA); rabbit polyclonal anti TNF-α (1:500, Novus Biologicals, Littleton, CO, USA); rabbit polyclonal anti IL-10 (1:200, Abbiotec, Escondido, CA, USA); mouse monoclonal anti CD68 (1:100, Bio-Rad, Hercules, CA, USA).

For fluorescence visualization of TH, IBA1 and CD68 a two-step indirect labelling protocol was used, while a three-step detection was performed to increase the signal of pSer129-αSyn, TNF-α and IL-10 by combining biotin-SP-conjugated IgG (1:500, Jackson Immunoresearch, West Grove, PA, USA) and streptavidin–fluorescein (1:400, Jackson Immunoresearch, West Grove, PA, USA), as previously described [44]. The sections were then incubated for 5□min in DAPI diluted in PBS before being mounted in Vectashield Antifade Mounting Medium (H-1000-10, Vector Laboratories, UK). Images were acquired using a spinning disk confocal microscope (Crisel Instruments, Rome, Italy) with a□×□63 magnification.

### TH neuronal and IBA^+^ microglial cell counting

The estimation of TH^+^ neurons and IBA1□ microglial cells was performed using ImageJ software (NIH, Bethesda, MD, USA) on images acquired with the Axio Scan.Z1 slide scanner (Zeiss, Germany) at 40× magnification. For TH neuronal counting, six sets of six midbrain sections, with 240 µm intersection interval, were acquired and analyzed for each animal in order to obtain a comprehensive representation of the area of interest. After background subtraction, TH^+^ neurons were manually counted rostro-caudally through the SNpc in sequential sections.

For microglial cell quantification, images were acquired using the DAPI channel to visualize nuclei and the RFP channel to detect IBA1□ cells; images were calibrated by setting the spatial scale using a pixel distance corresponding to 1000 µm, the SNpc was manually outlined, and the ROI area was measured (µm²). Background was then subtracted to facilitate cell identification, and IBA1□ cells were manually counted within the SNpc ROIs using the ImageJ “Cell Counter” tool on both the right and left sides of the midbrain. Finally, counts were normalized to the ROI area and expressed as cell density (cells/mm²). Mean values were calculated for each experimental group for graphical representation.

### Confocal microscopy analysis

Confocal imaging was performed using a spinning disk confocal microscope (Crisel Instruments, Rome, Italy) for both qualitative and quantitative analyses. Colocalization and volumetric analysis were performed in Imaris 7.3 software (Bitplane AG, Zurich, Switzerland), generating surfaces for each channel using uniform thresholds.

For the ThS analysis in SH-SY5Y cells challenged with α-Syn PFFs and treated with POM or 3MP, confocal z-stacks were acquired using a 63× oil immersion objective with a z-step size of 0.1 µm and a total optical section thickness of 4 µm. ThS□ signal was quantified in Imaris by generating a 3D surface on the ThS channel using uniform threshold parameters and expressed as ThS□ volume per cell (µm³/cell) by normalizing the total ThS□ volume to the number of cells in the same field.

For the quantification of pSer129-αSyn within the dopaminergic compartment of the SNpc, 8-16 systematically selected regions of interest (ROIs) measuring 700 × 700 × 30 µm were acquired and analyzed per animal. A colocalization channel between TH and pSer129-αSyn was generated within each ROI, so that only phosphorylated α-Syn located within TH^+^ structures was included in the analysis. For each ROI, we quantified both the total TH^+^ volume and the volume of colocalized pSer129-αSyn signal within TH^+^ structures. The pSer129-αSyn/TH ratio was then calculated as the volume of colocalized pSer129-αSyn divided by the TH^+^ volume within the same ROI. Values were averaged across ROIs for each animal and used for statistical analysis.

Imaging parameters for these analyses included a 63× oil immersion objective, a z-step size of 0.5 µm, and a total optical section thickness of 30 µm. The same imaging parameters were applied for the analysis of pro-and anti-inflammatory cytokines (TNF-α, IL-10). Cytokine expression was assessed at the single-cell level in IBA1^+^ microglial cells in both the SNpc and ACC, by analyzing between 50 and 90 cells per experimental group. Specifically, using the same Imaris-based colocalization approach, the overlapping volume between cytokine and IBA1 surfaces was quantified using the built-in colocalization tool, and the colocalization ratio was calculated as the volume of cytokine signal within IBA1^+^ cells relative to the total cell volume.

#### 3D Image Analysis of Microglial α-Synuclein Compartmentalization

Triple immunofluorescence for IBA1, CD68, and pSer129-αSyn was analyzed in Imaris 7.3 software from SNpc Z-stacks acquired at 100x (z-step size 0.3 µm; 35 µm thickness). For each experimental group, 30-50 individual microglial cells were analyzed. Raw 3D stacks were first pre-processed in Fiji/ImageJ by background subtraction for each channel (rolling ball radius = 200 pixels), using identical settings across all images. The background-corrected stacks were then imported into Imaris for 3D analysis.

3D surfaces were generated in Imaris (Surface module), using identical settings across groups. For each channel (IBA1, CD68, pSer129-αSyn), a single absolute intensity threshold was selected and kept constant across all images to include clearly positive signal and exclude background and low-intensity noise, and enable volumetric quantification. The IBA1 channel was used to segment individual microglial cells, followed by 3D visual inspection to remove/correct merged or truncated objects. The resulting IBA1 surfaces defined total cell volume (V_cell_, µm³). CD68 signal was masked to the IBA1 surface to retain only intracellular phagolysosomal structures (adapted from [45]). CD68^+^ objects were generated within this mask applying a minimum object volume cutoff (>0.01 µm³) and their summed volume per cell defined the phagolysosomal compartment volume (V_phago_).

pSer129-αSyn was analyzed in two steps: (i) masking pSer129-αSyn to the IBA1 surface to obtain total intracellular pSer129-αSyn volume (V_pSyn,total_); (ii) masking pSer129-αSyn to the CD68□ compartment within each IBA1□ cell, to obtain phagolysosomal pSer129-αSyn volume (V_pSyn,phago_). The non-phagolysosomal pSer129-αSyn fraction was calculated as V_pSyn,cyto_ = V_pSyn,total_ – V_pSyn,phago_. Two indices normalized to V_cell_ were derived: PCI = V_pSyn,phago_/V_cell_ and CDI = V_pSyn,cyto_/V_cell_, representing pSer129-αSyn localized in CD68□ phagolysosomes versus dispersed in non-CD68□ microglial compartments, respectively.

### Statistics

All statistical analyses were performed using GraphPad Prism 7 (San Diego, CA, USA) by researchers blinded to the experimental conditions. Data are presented as mean ± SEM. The assumption of normality was assessed using the Shapiro–Wilk test, while homogeneity of variances was evaluated with the Bartlett’s test and Levene’s test in RStudio (R version 4.5.2, R Foundation for Statistical Computing, Vienna, Austria). The specific statistical tests used, and details of data presentation are described in the figure legends. A p-value of less than 0.05 was considered statistically significant.

## Results

### 3MP mitigates α-synuclein-induced toxicity and inflammation *in vitro*

α-Syn, when incubated for 72 hr with rat primary mesencephalic mixed cultures, reduced cell dopaminergic neuron viability by approximately 50% (Fig. 1B, evaluated by counting TH^+^ stained cells). 3MP mitigated αSyn mediated neurotoxicity in a concentration-dependent mode across 3 to 60 µM range (Fig. 1B). Moreover, α-Syn induced microglial cell activation, as evaluated by IBA1 intensity, and a functionally proinflammatory profile shown by increased TNF-α production (Fig. 1C, D). 3MP concentration-dependently reduced both α-Syn-mediated IBA1 intensity and TNF-α production (Fig. 1C, D).

**Figure 1.**
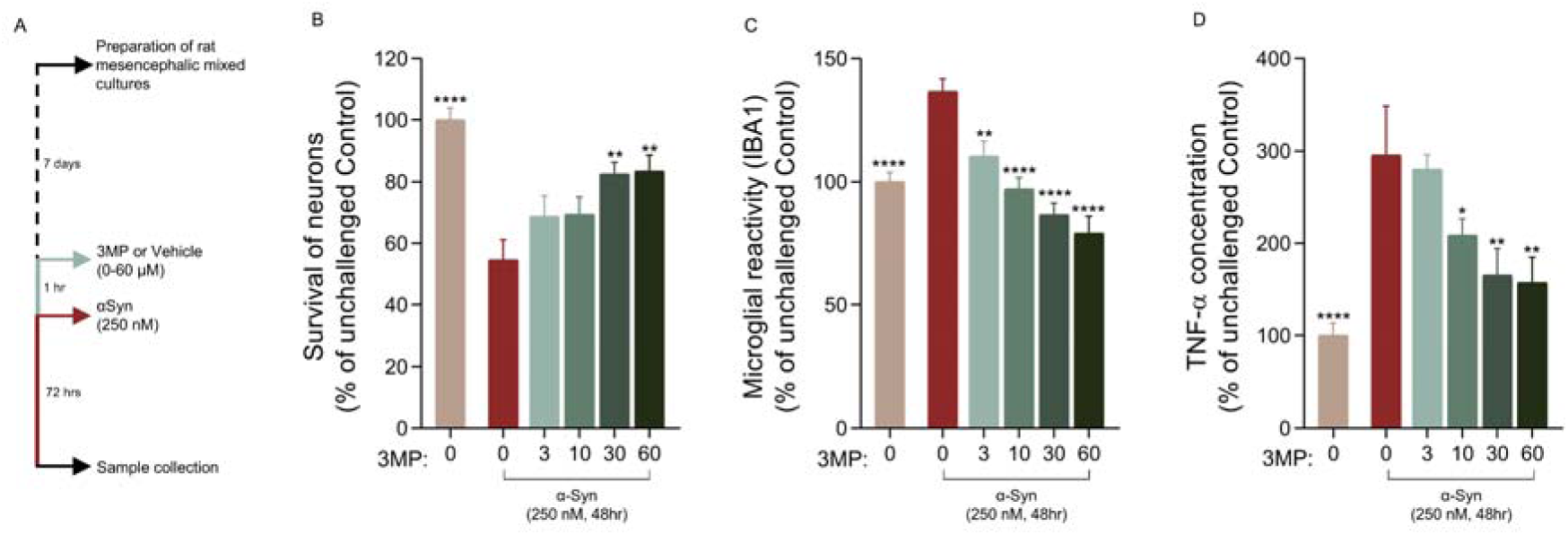
3MP ameliorated neuronal cell death and inflammation in rat primary mesencephalic mixed cultures challenged with αSynOs. (**A**) Experimental timeline for in vitro studies. (**B**) Mitigation of neuronal cell death, (**C**) microglial cell reactivity, and (**D**) TNF-α generation induced by αSyn (250 nM, 48 hr exposure). *p<0.05; **p<0.01; ***p<0.001; ****p<0.0001 vs. αSyn.

### 3MP Inhibits α-Synuclein Aggregation in Cell-Free Systems

We monitored the aggregation kinetics of α-Syn *in vitro* using a well-established assay to detect amyloid formation via ThT fluorescence. In this aggregation experiment, α-Syn aggregation was induced in PBS at 37 °C and in the presence of 2.5 mM CaClL and at a protein concentration of 100 μM. The measurements were performed under shaking and monitored for 120h. Under these conditions, ThT fluorescence rapidly increases and reaches a plateau after 72h (Fig. 2A). We confirmed the presence of fibrillar structures using transmission electron microscopy (TEM) imaging, revealing straight elongated α-Syn amyloid structures (Fig. 2D). When the incubation was performed with 1:1 molar ratio of α-Syn:POM, we observed a mild impairment of α-Syn aggregation, with the ThT plateau reaching 80% of that obtained in the absence of POM. The α-Syn fibrils grown in the presence of POM resulted generally shorter than the control, although showing a similar morphology. In addition, TEM images showed the presence of non-fibrillar amorphous aggregates in the presence of POM. When analysing the kinetics of aggregation, t ½ of values of α-Syn in the presence or absence of POM showed minimal (statistically non-significant) differences (Fig. 2B, C).

**Figure 2.**
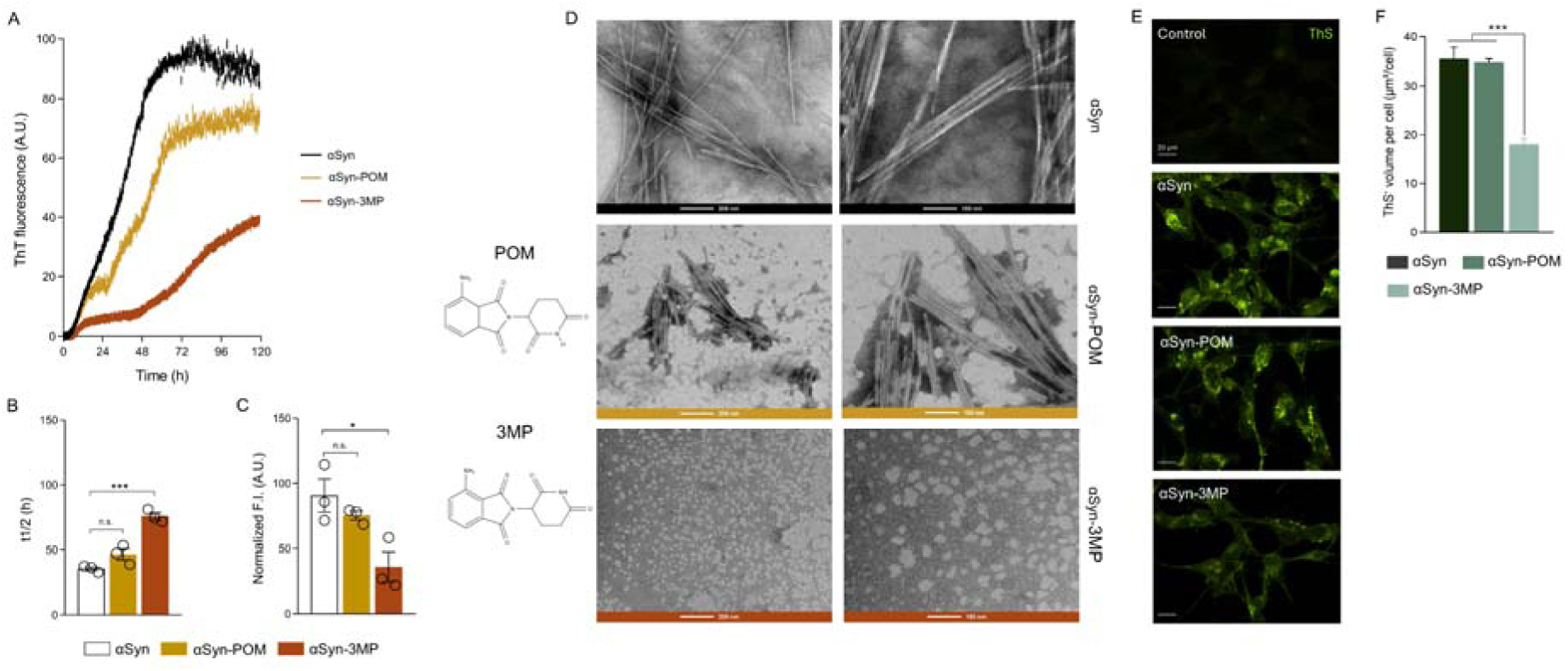
Effects of Pom and 3MP on α-Syn aggregation *in vitro*. (**A**) ThT fluorescence intensity monitored during α-Syn (100 μM) aggregation at 37 °C, in PBS at pH 7.4 and under orbital shaking (200 rpm). Kinetic traces for isolated α-Syn (black) and in co-incubation with POM (yellow) and 3MP (orange) are reported. All conditions were tested in triplicate. The intrinsic fluorescence of 3MP was found to be negligible compared to the intensity of ThT fluorescence. (**B**) t½ values of the kinetic curves. (**C**) Plateau values of the kinetic curves. One-way ANOVA followed by Dunnett’s post-hoc test was performed against protein alone (control). ns = not significant, * p < 0.05, *** p < 0.001. Shapiro–Wilk and Bartlett’s tests confirmed normality and homogeneity of variance. (**D**) Transmission electron microscopy (TEM) images of samples post 120h of incubations. The images were taken at both 100 nm and 200 nm magnifications. Samples were prepared using negative staining with 2% phosphotungstic acid (PTA) on carbon-coated 200 mesh copper grids. (**E**) Representative images of ThS staining on SH-SY5Y cells challenged with α-Syn pre-formed fibrils and treated with POM and 3MP (magnification 63x; scale bar: 20µm). (**F**) Quantification of ThS^+^ signal expressed as ThS□ volume per cell (µm³/cell) in SH-SY5Y cells challenged with α-Syn pre-formed fibrils and treated with POM or 3MP. Data are shown as mean ± SEM from three independent biological replicates. One-way ANOVA followed Tuckey’s post-hoc test.

A different scenario was found when incubating α-Syn with 1:1 molar equivalent of 3MP. In this case, α-Syn aggregation was found to be significantly inhibited, with ThT fluorescence curves being strongly affected in both the plateau (reduced to 40% of the control) and in the kinetics (with t ½ increasing from 36h in the control to 76h in the presence of 3MP) (Fig. 2B, C). Upon addition of POM or 3MP to pre-formed fibrils, ThT fluorescence was found to reach plateaus similar to those observed when incubation with these molecules was initiated from α-synuclein monomers (Fig. S1A). Moreover, TEM images of α-Syn aggregates in the presence of 3MP were significantly altered compared to the control, indicating a significant amount of amorphous assemblies as a result of the incubation in the presence of the small molecule (Fig. 2D). The size distribution of aggregated species formed in the presence of 3MP was further monitored by dynamic light scattering (DLS, Fig. S1B). The DLS profile obtained in the presence of 3MP revealed the emergence of a new population of aggregated α-synuclein species characterized by a significantly smaller size than mature fibrils. In addition, 3MP induced a concomitant reduction in the fibrillar population and an increase in the fraction of monomeric species.

### 3MP Prevents α-Synuclein Aggregation in a Cell-Based Model

α-Syn aggregation assay in SH-SY5Y cells was assessed *in vitro* using ThS staining following exposure to α-Syn PFFs. After four days, robust cytosolic ThS^+^ aggregates were observed, reflecting substantial intracellular α-Syn aggregation (Fig. 2E). Co-treatment with POM (αSyn-POM) failed to attenuate aggregates formation, as evidenced by persistent ThS signal comparable to the α-Syn group (Fig. 2E-F). In contrast, 3MP treatment (αSyn–3MP) markedly reduced ThSD inclusions (Fig. 2E) and significantly decreased the ThSD burden when quantified as ThSD volume per cell (µm³/cell) (Fig. 2F), supporting the efficacy of 3MP in limiting α-Syn aggregation in this cellular model. Control cells displayed minimal ThS fluorescence, indicating negligible baseline aggregation.

### 3MP is well tolerated in healthy rats

Before testing 3MP in the H-αSynOs–based PD model, we performed a pilot tolerability experiment in age-matched healthy rats to verify whether repeated dosing could affect general health status and baseline motor performance. Animals received 3MP intraperitoneally for two months either at 5 mg/kg on alternate days or at 10 mg/kg daily and were monitored longitudinally. Under these conditions, body weight gain remained comparable across groups, with no detectable deviation in the growth curve compared to Veh-Sal-treated animals (Fig. 3A). In addition, assessment of motor coordination using the challenging beam-walking test showed no changes in motor performance at all doses (Fig. 3B), supporting the absence of overt drug-related toxicity on somatic growth and gross motor behavior. Histology of the liver tissue showed that, at the macroscopic level, no gross hepatic lesions or signs of steatosis were observed in treated animals. Furthermore, histochemical analysis of the right medial lobe did not reveal inflammatory cell infiltration or evident structural damage following treatment compared to control animals. Hepatocytes displayed preserved cellular morphology, with no detectable alterations in cytoplasmic membrane or nuclear integrity (Fig. 3C).

**Figure 3.**
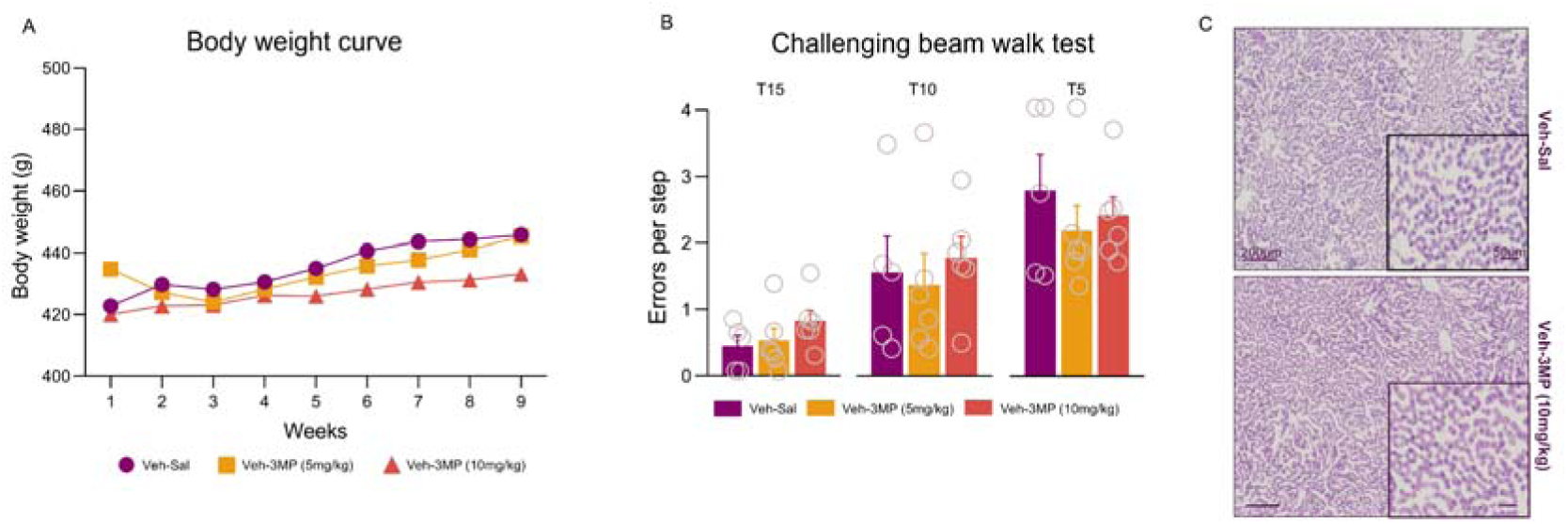
3MP tolerability in healthy rats. **(A)** Body weight curve of healthy rats receiving 5 or 10 mg/kg of 3MP intraperitoneally for two months and monitored longitudinally. (B) Errors per step committed on the three beams during the challenging beam walk test. (C) Representative H&E-stained micrographs of 5-µm coronal cryostat sections of the right medial liver lobe from paraformaldehyde-perfused rats infused with PBS and treated with saline or 3-MP (10 mg/kg). Scale bars: 200 µm; inset: 50 µm.

### 3MP rescues neuropathology and behavioral phenotype in a rat model of HαSynO-induced PD-like syndrome

We assessed the neuroprotective properties of 3MP in a previously validated model of PD obtained by the intranigral infusion of HαSynOs. 3MP was chronically administered starting at one-month post-HαSynOs infusion and for a two-months period. In previous studies we have shown that the one-month time point (post- HαSynOs) is associated with prodromal neuropathology, including presence of intraneuronal pSer129-αSyn aggregates, neuroinflammation and mitochondrial dysfunction, offering a useful window for therapeutic intervention.

#### Sensorimotor function

A challenging beam test was applied to assess motor performance and the impact of 3MP on motor impairment induced by exogenous HαSynOs infusion (Fig 4A). Consistent with previous findings [33,36], three months post-infusion HαSynOs-infused rats displayed a significantly higher number of errors per step compared to their Veh counterparts, across all three beams, indicating sensorimotor impairment (Fig. 4A). A two-way ANOVA for errors committed per step revealed a significant effect of infusion (F1,31 = 14.72, p<0.001) on the 15 mm beam, but no significant effect of treatment (F2,31 = 1.307, p≥0.05) or interaction between infusion and treatment (F2,31 = 2.419, p≥0.05). On the 5 mm beam, two-way ANOVA revealed a significant main effect of treatment (F2,28 = 9.958, p<0.001), whereas the main effect of infusion did not reach significance (F1,28 = 3.277, p≥0.05) and neither did the infusion × treatment interaction. (F2,28 = 2.720, p≥0.05). In contrast, on the 10 mm beam, significant effects were observed for both infusion (F1,29 = 16.67, p<0.001) and treatment (F2,29 = 5.751, p<0.01), as well as a significant interaction between infusion and treatment (F2,29 = 6.676, p<0.01). Importantly, chronic 3MP treatment, administered at two doses (5 and 10 mg/kg) and regimens, was associated with improved performance, most clearly on the 10 mm beam, where the significant infusion × treatment interaction indicates an infusion-dependent effect (Fig. 4A). Specifically, Tukey’s post-hoc comparisons showed a significantly lower number of errors per step in the HαSynOs group chronically treated with both the lower dose (5 mg/kg, every other day) and the higher dose (10 mg/kg, daily) on the 10 mm beam and also identified treatment-related reductions on the 5 mm beam.

**Figure 4.**
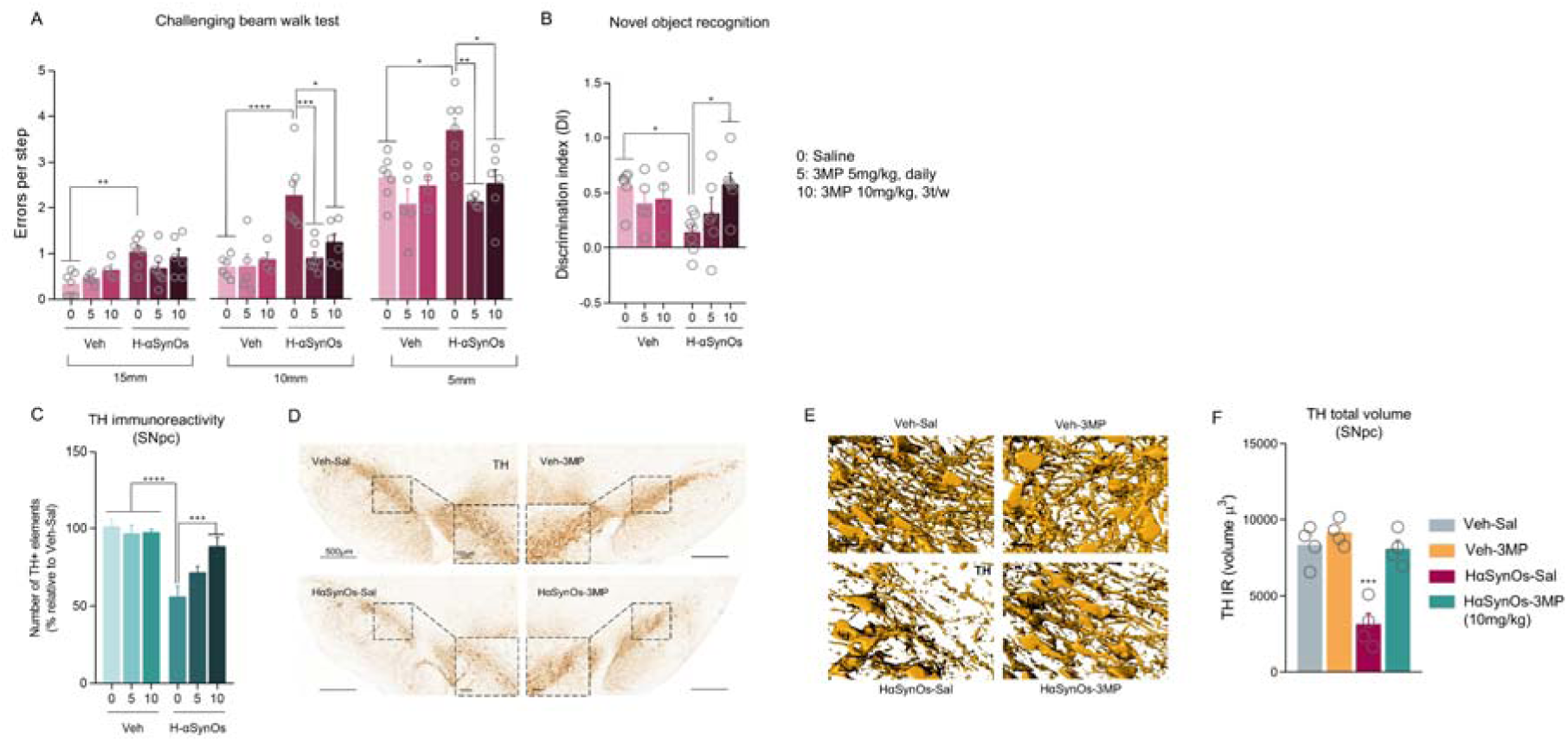
Symptomatologic and neuropathological PD-like traits were ameliorated by 3MP treatment. **(A)** Errors per step committed on the three beams during the challenging beam walk test. Data points represent individual animals (n = 4-7). *p<0.05, ** p<0.01, *** p<0.001, **** p<0.001 by two-way ANOVA followed by Tukey’s post hoc test. (**B**) Discrimination index measured by the novel object recognition test. Each dot represents individual animals (n = 4-7; * p < 0.05 by two-way ANOVA and Tukey’s post hoc test). **(C)** Number of TH^+^ elements measured along the SNpc. Values are the mean ± SEM; ***p<0.001, ****p<0.0001 by two-way ANOVA followed by Tukey’s post hoc test (n = 4-7). **(D)** Representative images of TH-stained SNpc slices (magnification 20×; scale bar: 500 μm). **(E)** 3D reconstruction of TH^+^ somas and fibers in the SNpc (magnification 63×; scale bar: 20 μm). **(F)** Total volume occupied by TH^+^ elements in the SNpc. Data were analyzed by two-way ANOVA followed by Tukey’s multiple comparisons test, with ***p < 0.001 indicating statistical significance. Each data point represents the average value from one animal, based on 8–16 ROIs per animal, with a total of n = 4 rats per experimental group.

#### Cognitive function

As cognitive disturbances are common non-motor symptoms in PD, we investigated whether the newly synthesized IMiD 3MP was also effective in mitigating cognitive impairments in parkinsonian rats with the short-term memory novel object recognition (NOR) task (Fig. 4B). Three months post-infusion, the HαSynOs-Sal group displayed a mild impairment in discriminating competence, as evidenced by a lower DI compared to Veh-infused rats (p<0.05), indicating a mild cognitive deficit (Fig. 4B). As shown in Fig. 4B, 3MP treatment was associated with an increase in DI within the HαSynOs condition, achieving a significant effect at the high dose (10 mg/kg, daily) (p<0.05).

#### Dopaminergic neurodegeneration

Three months post-surgery, HαSynOs-Sal rats exhibited a significant reduction in TH^+^ neurons, as quantified rostro-caudally in the SNpc, with an approximate 45% neuronal loss compared to Veh-infused animals (p<0.0001 by two-way ANOVA and Tukey’s post-hoc test) (Fig. 4C, D). Notably, 3MP treatment dose-dependently reversed the neuronal loss. We observed an approximately 70% neuronal survival with the lower dose (5mg/kg, every other day), and a nearly 90% survival with the 10 mg/kg daily treatment (Fig. 4C). Given the superior neuroprotective efficacy observed at 10 mg/kg, all subsequent mechanistic analyses were performed in the 10 mg/kg treatment group. The neuroprotective effect of 3MP against HαSynOs-induced neurodegeneration was further confirmed by volumetric analysis of TH^+^ elements in the SNpc (Fig. 4E, F). The total volume of TH^+^ neurons was markedly reduced in the HαSynOs-Sal group, whereas the 3MP treatment (10 mg/kg daily) fully counteracted this effect (p<0.001 by two-way ANOVA and Tukey’s post-hoc test). Interestingly, 3D reconstruction of both TH^+^ somas and fibers (Fig. 4E) revealed that HαSynOs impaired neuronal integrity of individual SNpc neurons, as evidenced by the presence of discontinuous TH^+^ fibers. 3MP effectively prevented this neuronal damage, preserving the integrity of the dopaminergic network within the SNpc.

#### pSer129-αSyn neuropathology

The presence of α-Syn intracellular inclusions is recognized as a neuropathological hallmark of α-synucleinopathies, including PD. In this context, levels of pSer129-αSyn, a marker of toxic α-Syn, were assessed in the SNpc. pSer129-αSyn was measured within TH^+^ neurons and IBA1^+^ cells. Three months after HαSynOs-infusion, we observed a significant increase in pSer129-αSyn/TH colocalization in the SNpc of HαSynOs-Sal rats compared to Veh controls (p < 0.5, Welch’s one-way ANOVA and Games–Howell’s post hoc test; see Fig. 5C). Immunohistochemical analysis revealed punctate staining of pSer129-αSyn within dopaminergic neurons, and 3D image reconstruction further demonstrated a high density of pSer129-αSyn puncta predominantly localized within the soma, with additional staining detectable along neuronal fibers (Fig. 5D). Notably, chronic treatment with 3MP markedly reduced pSer129-αSyn inclusions within TH^+^ neurons to levels comparable to those of Veh-treated animals (HαSynOs-3MP vs. HαSynOs-Sal, p < 0.1). To determine whether the greater anti-aggregation efficacy of 3MP compared with POM, as observed *in vitro* (Fig. 2), was also confirmed *in vivo*, we additionally analyzed the latter treatment. Unlike 3MP, rats chronically treated with POM did not show any significant reduction in pSer129-αSyn inclusions within TH^+^ neurons, confirming that this compound poorly targeted α-Syn aggregates directly (Fig. 5C) Of note, the direct statistical comparison between HαSynOs-3MP and HαSynOs-POM groups did not reveal a statistically significant difference, likely due to the wide data distribution in the HαSynOs-POM group (Additional file 1: Table S1 – Figure S2).

**Figure 5.**
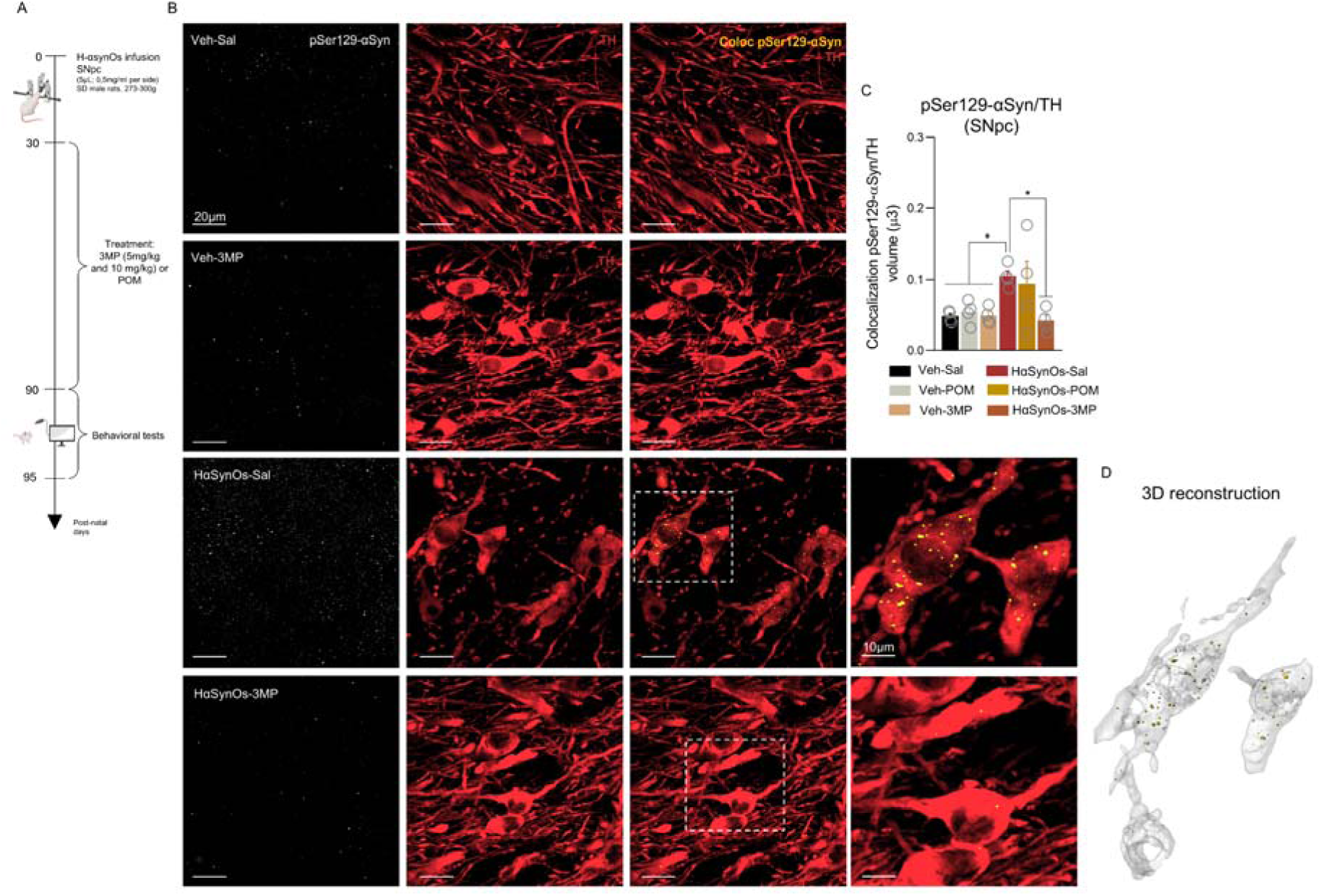
Pharmacological treatment with 3MP reduced the levels of toxic αSyn aggregates in mesencephalic dopaminergic neurons. (**A**) Experimental *in vivo* workflow (figure created with BioRender.com). (**B**) Representative immunostaining of total pSer129-αSyn (white) and colocalized pSer129-αSyn (yellow) within TH^+^ neurons (red). Images captured at 63x magnification; scale bar: 20 μm (first three columns), 10 μm (fourth column). (**C**) Volume of pSer129-αSyn colocalized within dopaminergic neurons. Values represent the mean ± SEM. Statistical analysis was performed using Welch’s one-way ANOVA with Games–Howell’s post hoc test to account for unequal variances; detailed pairwise comparisons are reported in Supplementary Table S1. p□value shown as *p□<□0.05. Each datapoint represents the average of 1 animal (8-16 ROIs per animal; with a total of n□=□4 rats per experimental group). (**D**) 3D reconstruction of pSer129-αSyn (yellow) aggregates within TH^+^ (white) neurons of HαSynOs-Sal treated rat.

Quantitative analysis of microglial cells in the SNpc revealed a trend towards an increased CD68/IBA1 volume ratio following HαSynOs infusion compared to the Veh counterparts, suggesting a possible expansion of the phagolysosomal compartment in response to α-Syn pathology (Fig. 6A), although differences did not reach statistical significance when assessed by the non-parametric Kruskal-Wallis test. Notably, the frequency distribution analysis of the CD68/IBA1 ratio for each experimental group revealed substantial differences in curve-shape (Additional file 1: Fig. S3). The HαSynOs-infused group displayed a clear bimodal distribution, indicative of the presence of two distinct subpopulations of responsive microglia with cells showing CD68 values comparable to controls, while others exhibiting a marked expansion of the phagolysosomal compartment, reflecting the underlying biological heterogeneity in response to α-Syn pathology. Interestingly, the PCI (Fig. 6B), representing the fraction of pSer129-αSyn localized in phagolysosomes within IBA^+^ microglial cells, showed an increase in HαSynOs-infused animals compared to controls (p<0.06 vs Veh-Sal; p<0.01 vs Veh-3MP). Similarly, the CDI (Fig. 6C), which reflects the cytoplasmic distribution of pSer129-αSyn, followed the same trend, being significantly elevated in HαSynOs-Sal animals, indicating enhanced accumulation of α-Syn aggregates in microglia under these conditions. Chronic treatment with 3MP normalized both the PCI and CDI in HαSynOs-infused animals, restoring these indices to control levels (p<0.01 vs Veh-Sal; p<0.0001 vs Veh-3MP and HαSynOs-3MP, Kruskal-Wallis test with Dunn’s post-hoc) (Fig. 6B-C). This effect is further corroborated by the 3D reconstructions shown in Fig. 5D, where IBA1^+^ microglia from HαSynOs-3MP animals display markedly reduced CD68 and pSer129-α-Syn immunoreactivity, closely resembling the morphology and staining patterns observed in the Veh-Sal group.

**Figure 6.**
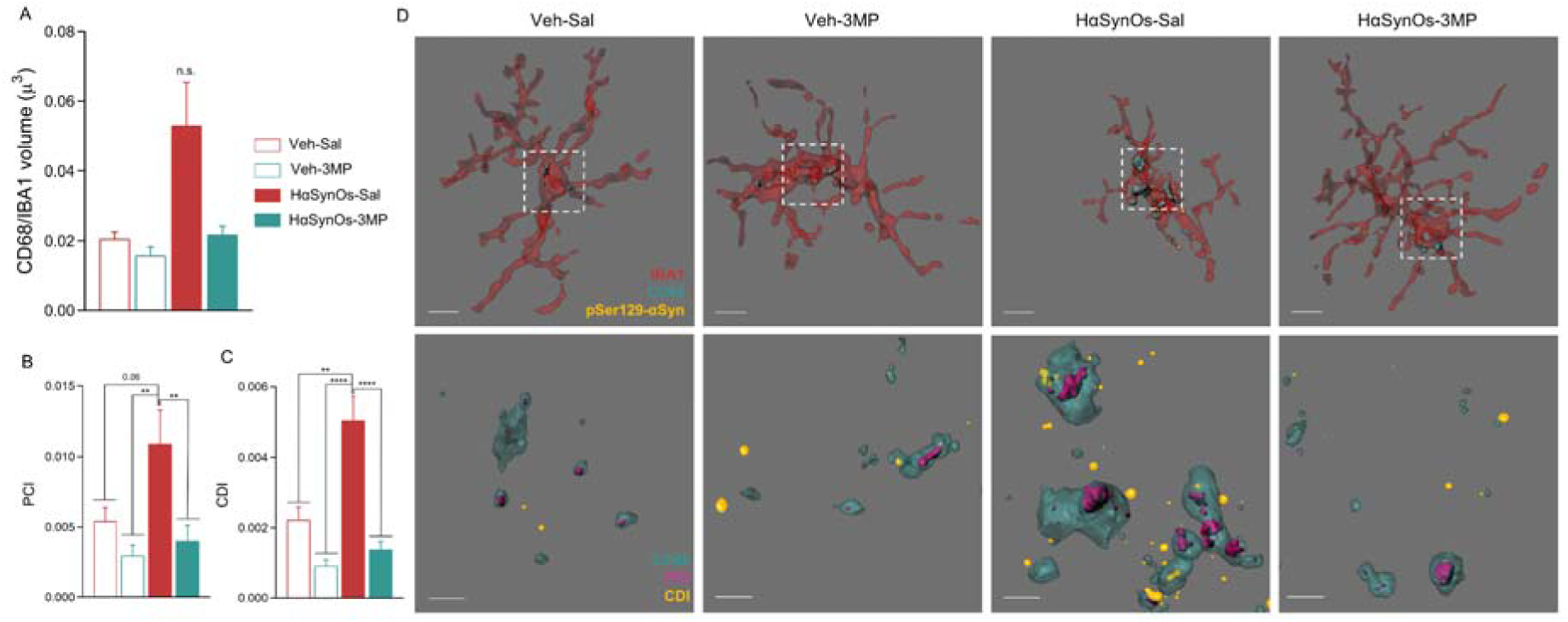
3MP treatment reduced pSer129-αSyn levels in microglial cells. **(A)** Quantification of the phagolysosomal compartment, expressed as the CD68/IBA1 volume ratio, in microglia from the SNpc across experimental groups. (**B, C**) Quantification of pSer129-αSyn aggregate distribution within IBA1^+^ microglia. The phagolysosomal colocalization index (PCI, **B**) and cytoplasmic distribution index (CDI, **C**) represent the fraction of pSer129-αSyn volume colocalized with CD68^+^ phagolysosomes or retained in the cytoplasm, respectively, normalized to IBA1 cell volume. Values are the mean ± SEM.**p < 0.01 and ****p < 0.0001, vs HαSynOs-Sal, Kruskal-Wallis with Dunn’s post hoc test (n = 30-50 cell per experimental group, n = 4 animals per group). (**D**) Representative 3D reconstructions of microglia (IBA1, red) in the SNpc from each experimental group, showing colocalization of CD68 (cyan) and pSer129-αSyn (yellow). Insets highlight the spatial distribution of aggregates within microglial cells. Lower panels display high-magnification views of the boxed regions, with CD68 (cyan), PCI (yellow), and CDI (magenta) channels separated for clarity. Magnification 100x. Scale bars: upper panels, 4 μm; lower panels, 1 μm. Veh-Sal: vehicle infusion into the SN, saline treatment; Veh-3MP: vehicle infusion into the SN, 3MP treatment (10 mg/kg 3t/w, i.p); HαSynOs-Sal: HαSynOs infusion into the SN, saline treatment; HαSynOs-3MP: HαSynOs infusion into the SN, 3MP treatment.

As previously reported [27], immunofluorescence did not detect aggregates of pSer129-αSyn in the ACC (data not shown).

#### Microglia immune response

IMiDs display the ability to modulate inflammatory responses by lowering TNF-α levels, which makes them attractive in the context of neuroinflammatory-related conditions. As previously reported [27,33], we first evaluated gross microglial reactivity in both the SNpc and ACC. Quantification of the total IBA1^+^ volume, did not reveal differences between HαSynOs-Sal and HαSynOs-3MP animals (Fig. 7A, B). While total IBA1^+^ volume was unchanged, 3MP decreased the number of IBA1^+^ microglial somata quantified along the rostro–caudal SNpc in HαSynOs rats (Fig. 7C). Consistently with the known immunomodulatory profile of IMiDs, Cytokine profiling revealed that 3MP significantly modulated the microglial intracellular production of TNF-α and IL-10 and their relative balance (Fig. 7D, E). Confirming previous studies [33], the HαSynOs infusion resulted in unbalanced intracellular production of the cytokines TNF-α and IL-10, with a marked increase in TNF-α levels alongside a decrease in IL-10 (Fig. 7D, E). Notably, 3MP was effective in reversing this ratio, strongly suppressing TNF-α production and boosting IL-10 levels in microglia (Fig. 7D, E). The analysis of TREM-2 in IBA1^+^ cells in the SNpc showed that the HαSynOs infusion induced a marked decrease in the expression, while 3MP treatment was associated with a modest upward trend toward control levels that did not reach statistical significance (Fig. 7F).

**Figure 7.**
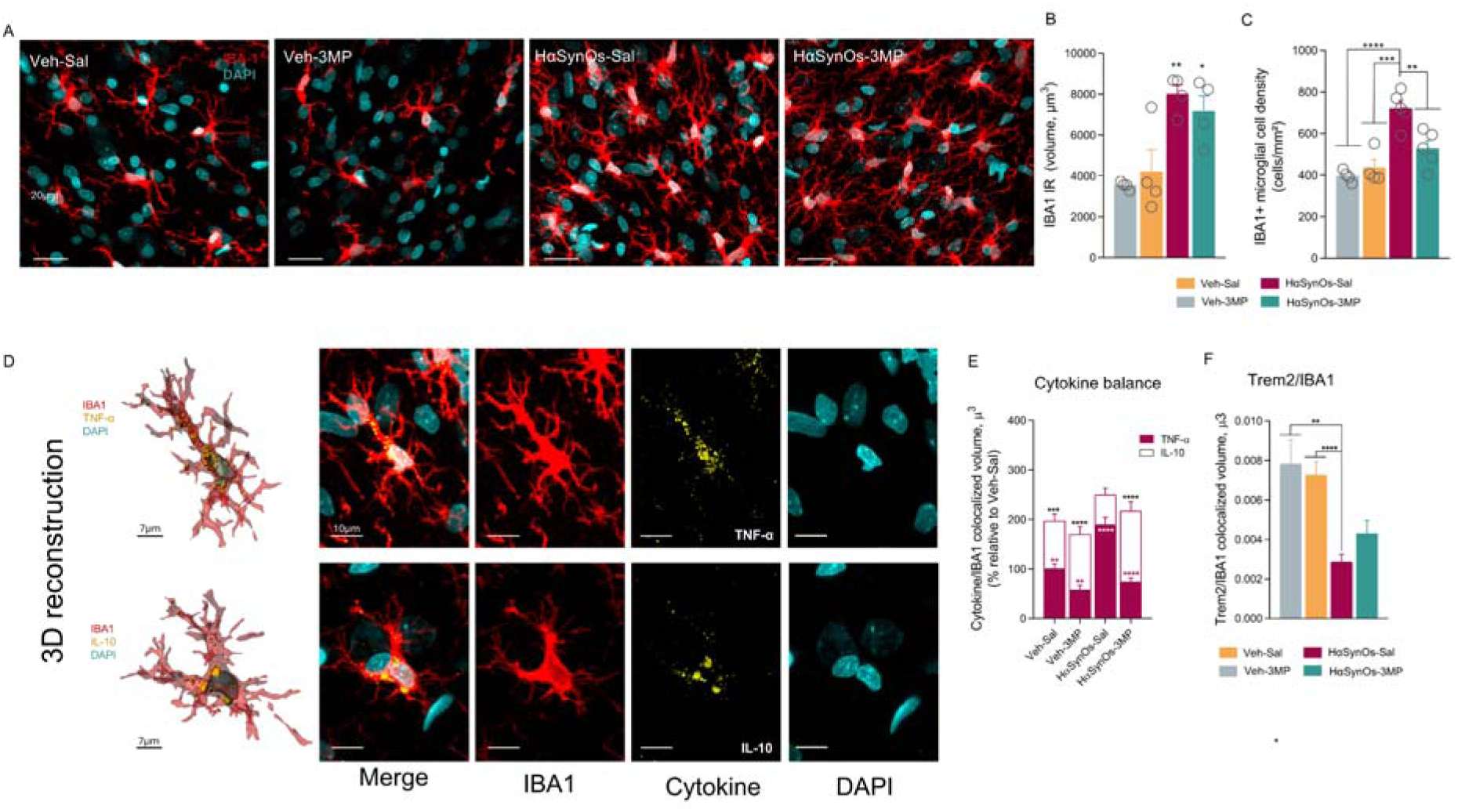
Midbrain microglial cells shifted their cytokine profile following 3MP treatment. (**A**) Representative immunostaining of microglial cells (IBA1, red) and nuclei (DAPI, blue) in the SNpc across four experimental conditions. Images were captured at 63x magnification; scale bar 20 μm. (**B**) Total volume occupied by IBA1^+^ cells in the SNpc. Data are presented as mean ± SEM. Statistical analysis was performed using two-way ANOVA with Tukey’s post hoc multiple comparisons test. Significance is indicated by *p < 0.05 (HαSynOs-3MP vs Veh groups) and p < 0.01 (HαSynOs-Sal vs Veh groups. Each data point corresponds to the average measurement from a single animal, calculated from 12–16 regions of interest (ROIs) per animal, with a total of n = 4 rats per experimental group. (**C**) IBA1^+^ microglial cell density in the SNpc (cells/mm²). Data are the mean ± SEM. Statistical analysis was performed using two-way ANOVA with Tukey’s post hoc multiple comparisons test. Each data point corresponds to the average measurement from a single animal (n = 4-5 rats per experimental group). (**D**) Percentages of the two colocalized cytokines TNF-α (magenta) and IL-10 (white) whithin IBA1□cells. Cytokines were analysed separately using the Kruskal–Wallis test, followed by Dunn’s multiple comparisons test. Magenta asterisks indicate significant differences in TNF-α intracellular production relative to HαSynOs-Sal; black asterisks indicate significant differences in IL-10. Within each experimental group, TNF and IL-10 percentages were compared using the Mann–Whitney U test (white asterisk). TNF expression was quantified from 50 cells per group; IL-10 from 60 cells per group (from 4 animals per group). (**E**) Trem2/IBA1 colocalized volume in the SNpc. The volume of Trem2 signal colocalized within IBA1□ microglia (µm³) was quantified across the four experimental groups. Data are shown as mean ± SEM. Statistical analysis was performed using the Kruskal–Wallis test, followed by Dunn’s multiple comparisons test. Measurements were obtained at the single-cell level (n = 60 microglial cells per group, sampled from 4 rats per group. Veh-Sal: vehicle infusion into the SN, saline treatment; Veh-3MP: vehicle infusion into the SN, 3MP treatment (10 mg/kg 3t/w, i.p); HαSynOs-Sal: HαSynOs infusion into the SN, saline treatment; HαSynOs-3MP: HαSynOs infusion into the SN, 3MP treatment.

Similarly, in the ACC, 3MP demonstrated a modulatory effect on the microglial response, reducing the intracellular production of TNF-α production while increasing IL-10 levels (Fig. 8A-D), thereby confirming the compound’s ability to rebalance inflammatory responses.

**Figure 8.**
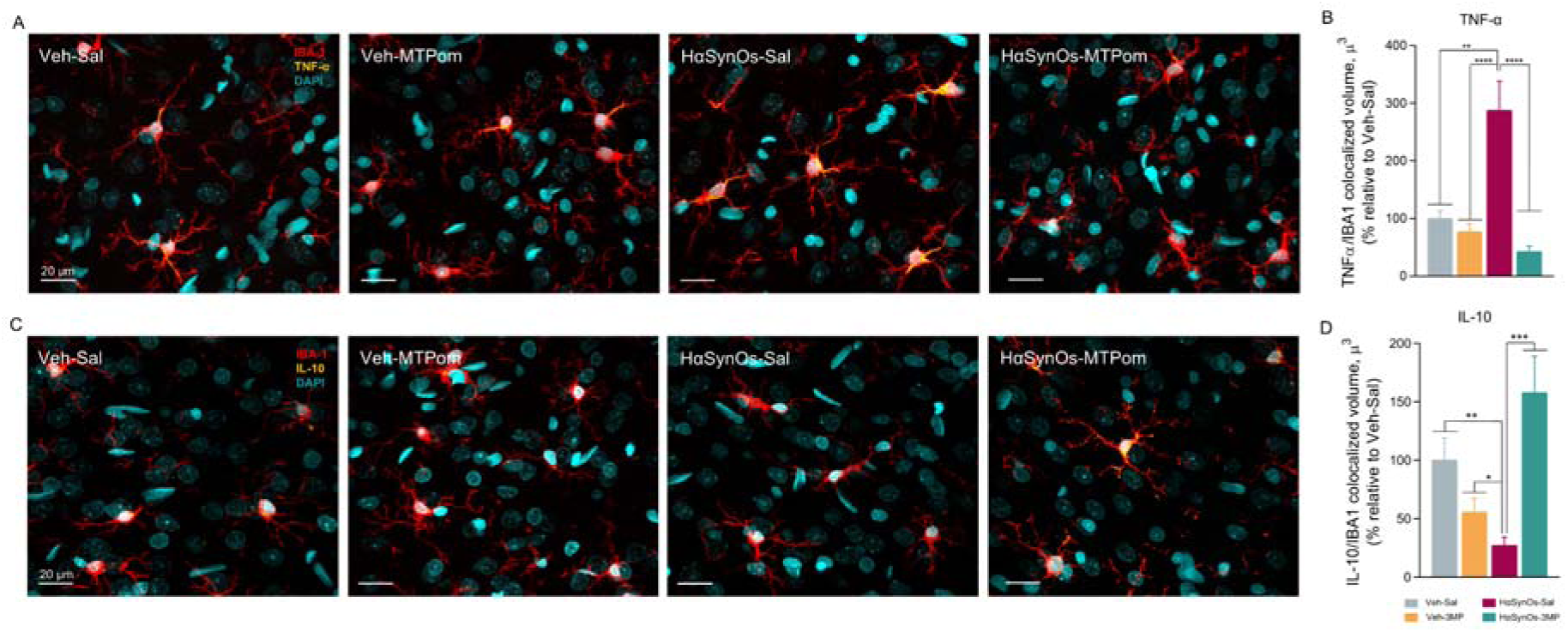
Treatment with 3MP restored the cytokine content in cortical microglial cells induced by the intranigral infusion of HαSynOs. (**A, C**) Representative images of colocalized TNF-α (yellow, upper) and IL-10 (yellow, lower) within IBA1^+^ cells (red). Magnification 63x, scale bar: 20 μm. Bar plot showing the volume of TNF-α (**B**) and IL-10 (**C**) colocalized within microglial cells in the ACC. Kruskal-Wallis and Dunn’s multiple comparison test. P□values shown as not significant *P□<□0.05; **P□<□0.01; ***P□<□0.001; ****P□<□0.0001 (n = 70-100 cells per group, n = 4-5 animals per group).

## Discussion

In the present study, by using a combined *in vitro* and *in vivo* approach, we demonstrated that the novel IMiD 3MP effectively mitigated PD-like pathology and provided first evidence of improved motor and cognitive readouts induced by toxic α-SynOs, by simultaneously targeting the α-Syn neuropathology and accompanying neuroinflammation, thereby positioning 3MP as a potential disease-modifying agent for PD. The newly synthesized IMiD, 3MP, may offer an alternative to traditional IMiDs, such as POM. Whereas POM, a third-generation thalidomide analogue, has demonstrated efficacy across diverse neurological conditions [30,33,46], it shares known limitations of the IMiD class, including teratogenicity and potential systemic toxicity. Such teratogenic action is, in large part, mediated by interaction with CRBN, a substrate receptor for the CRL4^CRBN^ E3 ubiquitin ligase complex [47]. Whereas CRBN binding underpins the useful anti-myeloma activity of this class of compounds by modifying the neosubstrate preference degraded by the E3 ligase [48], this same CRBN interaction leads to degradation of the neosubstrate Sall4, a transcription factor critical for normal embryonic development [49,50]. In contrast, 3MP potently binds to CRBN without inducing, alike, rapid Sall4 degradation [34], potentially lowering teratogenic risk, as recently evaluated by an absence of toxicity in chicken embryo assays [34].

This different mechanism of action, evaluated in molecular modelling studies of differing 3MP and POM chemical interactions with human CRBN [34], highlights 3MP as a compelling IMiD candidate for the treatment of neurological disorders, for which a more potent and potentially safer IMiD could be valuable [51].

As part of our initial screening, we assessed the efficacy of 3MP in rat primary mesencephalic mixed cultures challenged with α-Syn and subsequently treated with escalating 3MP doses. Notably, 3MP effectively reduced neuronal cell loss and displayed a potent and dose-dependent anti-inflammatory effect, mitigating the inflammatory cascade triggered by α-Syn challenge. Such anti-inflammatory action aligns with the ability of 3MP to mitigate lipopolysaccharide (LPS)-induced inflammation in RAW 264.7 (macrophage) and IMG (microglial) cells and reduce systemic and brain inflammatory markers in LPS challenged rodents [32,34]. The direct anti-aggregation properties of 3MP were assessed by the ThT *in vitro* aggregation assay, a chemical method that evaluates the formation of α-Syn fibrillary species independently of cellular systems. This approach allowed us to directly compare the effects of 3MP and POM on the aggregation of α-Syn monomers. Notably, 3MP, in contrast to POM, effectively inhibited α-Syn fibrillization, suggesting a specific molecular mechanism that may underlie its therapeutic efficacy in reducing α-Syn pathology. Thereafter, to validate these findings in a more physiologically relevant context, we performed the ThS assay in cellular model. SH-SY5Y cell cultures treated with either 3MP or POM and subsequently exposed to α-Syn, were assessed for α-Syn aggregation using ThS fluorescence. Consistent with the acellular *in vitro* assay, only 3MP treatment resulted in a significant reduction of the ThS fluorescence signal, indicating its ability to directly interfere with α-Syn aggregation within a cellular environment.

Building on *in vitro* studies, the impact of 3MP on PD-like pathology was validated in a male rodent model of synucleinopathy, achieved through the bilateral intranigral infusion of H-αSynOs [33,36]. As an initial tolerability assessment, a prolonged treatment with escalating doses of 3MP in healthy rats did not alter the body weight curve, nor induced any modification of the motor performance. Moreover, the macroscopic analysis of the liver, evaluated as the main metabolizing organ for iMIDs, showed absence of gross hepatic lesions with preserved cell morphology, no signs of steatosis or inflammatory cell infiltration in both saline and 3MP-treated rats. Moreover, pharmacokinetic parameters, including brain and plasma concentration of 3MP administered as a nanosuspension were carefully evaluated in a separate study, showing sustained and elevated brain levels relative to plasma [35]. In line with this, in a recent study evaluating the efficacy of 3-MP in a rat model of ischemic stroke, 3-MP demonstrated an oral bioavailability of 38.5%, a brain/plasma concentration ratio of 0.5 to 0.6, and a plasma half-life compatible with once daily dosing [32]. In the PD model, we initiated the chronic pharmacological treatment with 3MP one month after the infusion of HαSynOs, a phase considered prodromal in our model and characterized with early signs of neuropathology. The choice of this early intervention underscores the translational relevance of both the disease model and the therapeutic strategy, emphasizing their potential to bridge preclinical findings with clinical applications. This timing aligns with previous findings showing that at this early staging, despite the absence of dopaminergic degeneration, there was a significant accumulation of pSer129-α-Syn aggregates in the SNpc confined to dopaminergic neurons, invading microglial cells later as neuropathology progressed alongside a robust inflammatory response [36].

In line with our previous observation with POM [33], 3MP significantly modulated intracellular cytokine production by microglial cells, in accord with its immunomodulatory properties and well-known mechanism of action of iMIDs, which target TNF-α synthesis [47]. Specifically, infusion of HαSynOs resulted in a dysregulated cytokine profile in microglia, characterized by elevated TNF-α and decreased IL-10 levels, which was effectively reversed by 3MP. This shift is particularly relevant given TNF-α’s role in exacerbating neuroinflammation and neuronal damage, and IL-10 in reparative processes.

As a novel and compound-specific property, providing *in vivo* confirmation of the results initially observed in the *in vitro* models, we found that 3MP significantly reduced protein aggregates within neurons and in microglia during a two-months treatment, an effect paralleled by a significant protection of dopaminergic neurons of the SNpc. To confirm the improved profile of 3MP versus POM revealed by *in vitro* ThT assays, we evaluated pSer129-α-Syn accumulation in the SNpc of rats treated with POM for two months in the same PD model [33]. The cumulative frequency profiles of pSer129-α-Syn revealed that, unlike 3MP that was associated with a consistent reduction of intraneuronal aggregates across all animals, POM yielded a variable response, with high-burden cases persisting within the treated group. Together, these observations are compatible with a divergent *in vivo* activity profile of the two IMiDs and a qualitative improvement of 3MP over POM, potentially reflecting differences in target engagement and/or downstream mechanisms relevant to α-Syn aggregation and clearance.

Although the present data do not dissect mechanistic aspects, which are currently under investigation, we speculate that the lower presence of intracellular pSer129-αSyn aggregates induced by 3MP, may result either from an early neuronal drug-targeting, that would prevent intraneuronal formation and release of pSer129-α-Syn and the subsequent microglia uptake and overload, or from a direct effect on both cell populations, directly impacting clearance mechanisms of microglia. Notably, and regardless of the underlying mechanism, 3MP sustained a more physiological phagolysosome compartment, the cell structure mainly involved in the degradation of engulfed material in microglia. By investigating the compartmentalization of pSer129-α-Syn within microglia, we found a significant increase of pSer129-αSyn within the cytoplasmic compartment, defined as CDI, after the HαSynOs infusion. In addition, the volume of the CD68^+^ phagolysosomal compartment (CPI) was enlarged in this condition, containing abundant pSer129-αSyn aggregates. Of note, this group displayed a bimodal distribution, which reflects the presence of two distinct subpopulations of responsive microglia with one exhibiting a marked expansion of the phagolysosomal compartment, likely reflecting the underlying biological heterogeneity in response to α-Syn pathology. In contrast, 3MP treatment normalized the size of the phagolysosome and reduced the presence of pSer129-αSyn in both the phagolysosome and cytoplasmic compartments. Moreover, in this group the data distribution shape overlapped that of Veh-treated animals. Although these results should be interpreted as descriptive indices of compartmentalized intracellular α-Syn, being derived from a static 3D reconstruction at a single time point, they may reflect a HαSynOs-induced pathological accumulation in the phagolysosome and perhaps defective degradation, which is resolved by 3MP, aligning with previous findings [52] and our previous study showing that pre-exposure of microglia to H-αSynOs significantly reduced microglial phagocytic capacity [36]. In contrast, 3MP treatment normalized the size of the phagolysosome and reduced the presence of pSer129-αSyn in both the phagolysosome and cytoplasmic compartments. Of note, this was associated with a partial rescue of TREM-2 expression, a recognized marker of disease-associated microglia (DAM) [53]. In fact, in line with previous findings in neurodegenerative conditions including PD, and with a neuroprotective function attributed to this surface receptor, TREM-2 expression was significantly dampened after the H-αSynOs infusion in PD rats and partially restored by 3MP [54]. Parenthetically, the recent unexpected failure of the INVOKE-2 trial with a monoclonal antibody that targets TREM-2 has questioned the appropriateness of directly targeting TREM-2 expression in neurodegenerative diseases, suggesting context-dependent protective or detrimental actions of TREM-2 [55]. In this context, targeting upstream mechanisms which in turn restore physiological TREM-2 levels may provide an attractive therapeutic approach.

Altogether, findings from the *in vivo* model, together with the results obtained by *in vitro* assays, support the concept that early intervention with 3MP may not only target neuronal pathology but also attenuate the spread of aggregates from neurons to microglial cells [6,52], with broader implications for disease progression [54–56]. Early 3MP intervention may halt the neuron-to-glia propagation of α-Syn, prevent pathological microglia and neuroinflammation, and overall the microglial involvement in disease progression [59,60], reiterating the critical issue of early intervention in PD.

Notably, 3MP effectively countered neuronal death induced by toxic oligomeric α-Syn in a dose-dependent manner in male rats, achieving up to 90% neuronal survival. The 3D reconstruction of nigral dopaminergic neurons highlighted the significant disruption of the dopaminergic network, as shown by the reduced number of fibers and neuronal somata, as well as neurites in a state of discontinuity. This is consistent with the known effects of toxic oligomeric species, which compromise membrane integrity through their insertion into the phospholipid bilayer [4], and previous studies demonstrating that α-SynOs can severely disrupt neurite morphology in human dopaminergic neuronal cell lines [61]. Remarkably, pharmacological treatment with 3MP not only counteracted neuronal death induced by H-αSynOs but also preserved the morphological integrity of the dopaminergic network, mitigating the disruption of neurite morphology and maintained the structural coherence of the network.

The neuroprotective effects of 3MP observed at cellular and morphological levels were further corroborated by behavioral assessments in our model. The challenging beam walk test, a sensitive measure of motor deficits associated with nigrostriatal pathway degeneration, was used to evaluate motor function recovery [36,41,42]. Confirming previous results [33,36], the infusion of H-αSynOs significantly increased errors during the test, reflecting impaired motor coordination. Notably 3MP was associated with a reduction of errors, most clearly on the intermediate 10-mm beam, aligning with the neuroprotective effects of 3MP on the nigrostriatal circuit, and in line with previously shown IMiDs properties [33]. By contrast, the 15-mm and 5-mm beams may be less sensitive to detect treatment-related differences due to task difficulty, with the 15-mm beam being relatively easy (ceiling effect) and the 5-mm beam being highly challenging (floor effect), potentially diluting group differences compared with the intermediate condition [62].

Our study further expands our understanding of cognitive dysfunction in PD by providing evidence that chronic 3MP treatment modulates cognitive performance. Cognitive impairments, that range from MCI to PDD, are significant components of the clinical spectrum of PD, and interventions capable of mitigating these symptoms may significantly improve the quality of life for affected individuals [20]. Building on our previous findings [27,63], which identified H-αSynOs-induced dysfunction and neuroinflammation in the ACC as key contributors to cognitive deficits [27,63], we found here that 3MP was associated with an improvement in NOR performance, alongside reduced neuroinflammation in the ACC, suggesting its potential to counteract impaired cognition. 3MP improved the performance in the NOR test and restored microglial TNF-α/IL-10 ratio in ACC, a critical aspect for preserving neuronal function and supporting cognitive processes.

Altogether, our findings support the potential for disease modification in our PD models, highlighting a dual mechanism of action for 3MP, which includes a reduction of neuronal aggregates and modulation of microglial activity. These interconnected processes result in preserved neuronal integrity while enabling microglia to adopt a phenotype to resolve inflammation and support tissue repair. As a caveat and basis for next studies, it should be highlighted that the present *in vivo* experiments were conducted exclusively in male rats, since the H-αSynOs-induced PD model has so far been fully characterized only in males. Nevertheless, preliminary evidence of 3MP efficacy across sexes has recently been obtained in Drosophila, were 3MP was neuroprotective in both males and females, in two different PD models (Mocci et al, in preparation). Considering this, further studies are warranted to definitively ascertain the efficacy of IMiDs, and particularly of 3MP, in female mammalian preclinical models of PD.

## Conclusion

Our study supports the multifaceted therapeutic potential of 3MP in addressing key pathological hallmarks and provides first evidence of benefits on both motor and non-motor readouts in PD. Unlike many current pharmacological treatments for PD, which are often mono-targeted and primarily address symptomatic relief, 3MP demonstrates a unique capability to modulate multiple key pathological processes of PD, including α-Syn aggregation, neuroinflammation, and is associated with functional improvement in this preclinical model, to potentially provide a more holistic approach to the complexities of the disease.

## Abbreviations

3MP: 3-monothiopomalidomide
α-Syn: α-Synuclein
ACC: Anterior Cingulate Cortex
CRBN: Cereblon
H-αSynOs: Human α-Syn Oligomers
IL-10: Interleukin 10
IMiDs: Immunomodulatory Imide Drugs
LB: Lewy Bodies
MCI: Mild Cognitive Impairment
PD: Parkinson’s Disease
POM: Pomalidomide
TH: Tyrosine Hydroxylase
ThT: Thioflavin T
TNF: Tumour Necrosis Factor

## Declarations

### Ethics approval and Consent to participate

All experimental procedures were conducted in compliance with the ARRIVE guidelines and the European Community Directive 2010/63/EU (L 276, 20/10/2010). The study was approved by the Italian Ministry of Health (authorization no. 766/2020-PR).

### Consent for publication

Not applicable.

### Availability of data and materials

The datasets related to this study are available from the corresponding author on reasonable request.

### Competing interests

The authors declare that they have no competing interests.

### Funding

This study was supported by the Intramural Research Program, National Institute on Aging, NIH, USA (AG000994); MNESYS Spoke 7 (Neuroimmunology and Neuroinflammation): PROFILES, Funded by EU - Next Generation EU, Missione 4 Componente 2 investimento 1.3 (B33C22001060002). PRIN 2022: P20222F4JY; Aevis Bio, Inc., Republic of Korea (the Korea Dementia Research Center (KDRC), funded by the Ministry of Health & Welfare and Ministry of Science and ICT, Republic of Korea (RS-2024-0034124913); the Korea Drug Development Fund funded by the Ministry of Science and ICT, Ministry of Trade, Industry, and Energy, and Ministry of Health and Welfare, Republic of Korea (KDDF RS-2024-00338017).

### Author contributions

MFP was the principal investigator, responsible for experiments execution, data analysis and manuscript draft; KA, ME, JM contributed to behavioral experiments, immunohistochemistry and image analysis, MR and ADS prepared alpha-synuclein oligomers and performed the *in vitro* ThT assay; PP conducted *in vitro* studies for 3MP neuroprotecrive and anti-inflammatory assay; CP and VS were responsible for the aggregation assay in cell system; DT was in charge of drug synthesis, LC, MCC, FL prepared the nanoformulations of 3MP and POM; AP collaborated in rodents surgeries and behavioral testing; DSK. NHG contributed with drug synthesis, conceptualization and manuscript drafting, funds; ARC is the corresponding author, contributed to conceptualization, manuscript drafting, funding.

## Acknowledgements

We acknowledge Cesast (Academic Services Center for Animal Facility) for animal care and housing and Cesar (Academic Services Center for Research Support) for technical assistance at the University of Cagliari. We also acknowledge Dr. Gessica Piras for her skilled assistance throughout the *in vivo* experimentation, and Prof. Marta Kowalik for support in the examination of liver tissues for pathological assessment.

## Notes

### Competing Interest Statement

The authors have declared no competing interest.

