## Supplementary Figures and Table for "A new therapeutic approach for Parkinson’s disease: dual targeting of α-Synuclein aggregation and microglial function by the novel immunomodulator 3-Monothiopomalidomide"

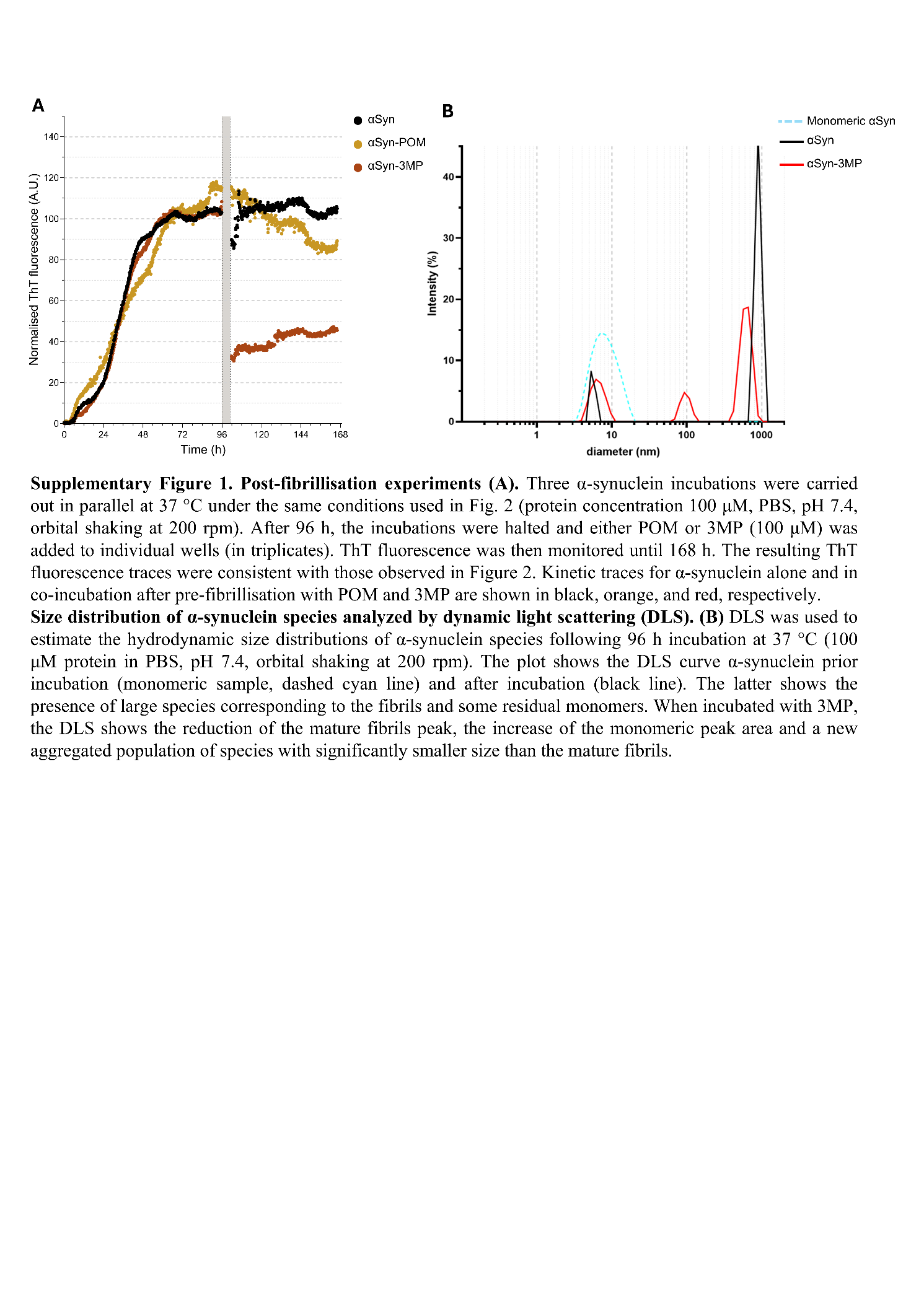

**Supplementary Figure 1. Post-fibrillisation experiments (A).** Three α-synuclein incubations were carried out in parallel at 37 °C under the same conditions used in Fig. 2 (protein concentration 100 μM, PBS, pH 7.4, orbital shaking at 200 rpm). After 96 h, the incubations were halted and either POM or 3MP (100 μM) was added to individual wells (in triplicates). ThT fluorescence was then monitored until 168 h. The resulting ThT fluorescence traces were consistent with those observed in Figure 2. Kinetic traces for α-synuclein alone and in co-incubation after pre-fibrillisation with POM and 3MP are shown in black, orange, and red, respectively.

**Size distribution of α-synuclein species analyzed by dynamic light scattering (DLS). (B)** DLS was used to estimate the hydrodynamic size distributions of α-synuclein species following 96 h incubation at 37 °C (100 μM protein in PBS, pH 7.4, orbital shaking at 200 rpm). The plot shows the DLS curve α-synuclein prior incubation (monomeric sample, dashed cyan line) and after incubation (black line). The latter shows the presence of large species corresponding to the fibrils and some residual monomers. When incubated with 3MP, the DLS shows the reduction of the mature fibrils peak, the increase of the monomeric peak area and a new aggregated population of species with significantly smaller size than the mature fibrils.

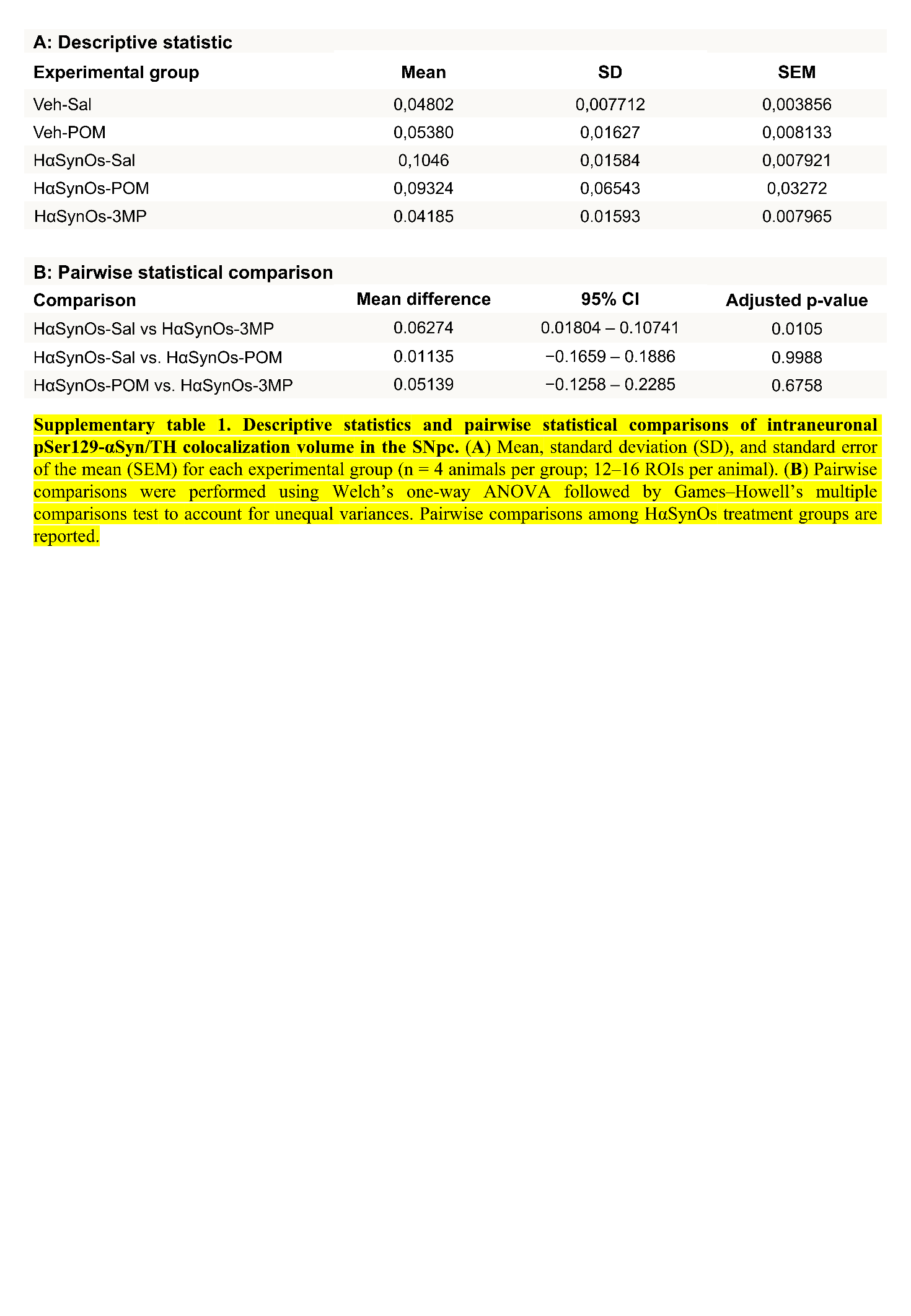

**Supplementary table 1. Descriptive statistics and pairwise statistical comparisons of intraneuronal pSer129-αSyn/TH colocalization volume in the SNpc.** (**A**) Mean, standard deviation (SD), and standard error of the mean (SEM) for each experimental group (n = 4 animals per group; 12–16 ROIs per animal). (**B**) Pairwise comparisons were performed using Welch’s one-way ANOVA followed by Games–Howell’s multiple comparisons test to account for unequal variances. Pairwise comparisons among HαSynOs treatment groups are reported.

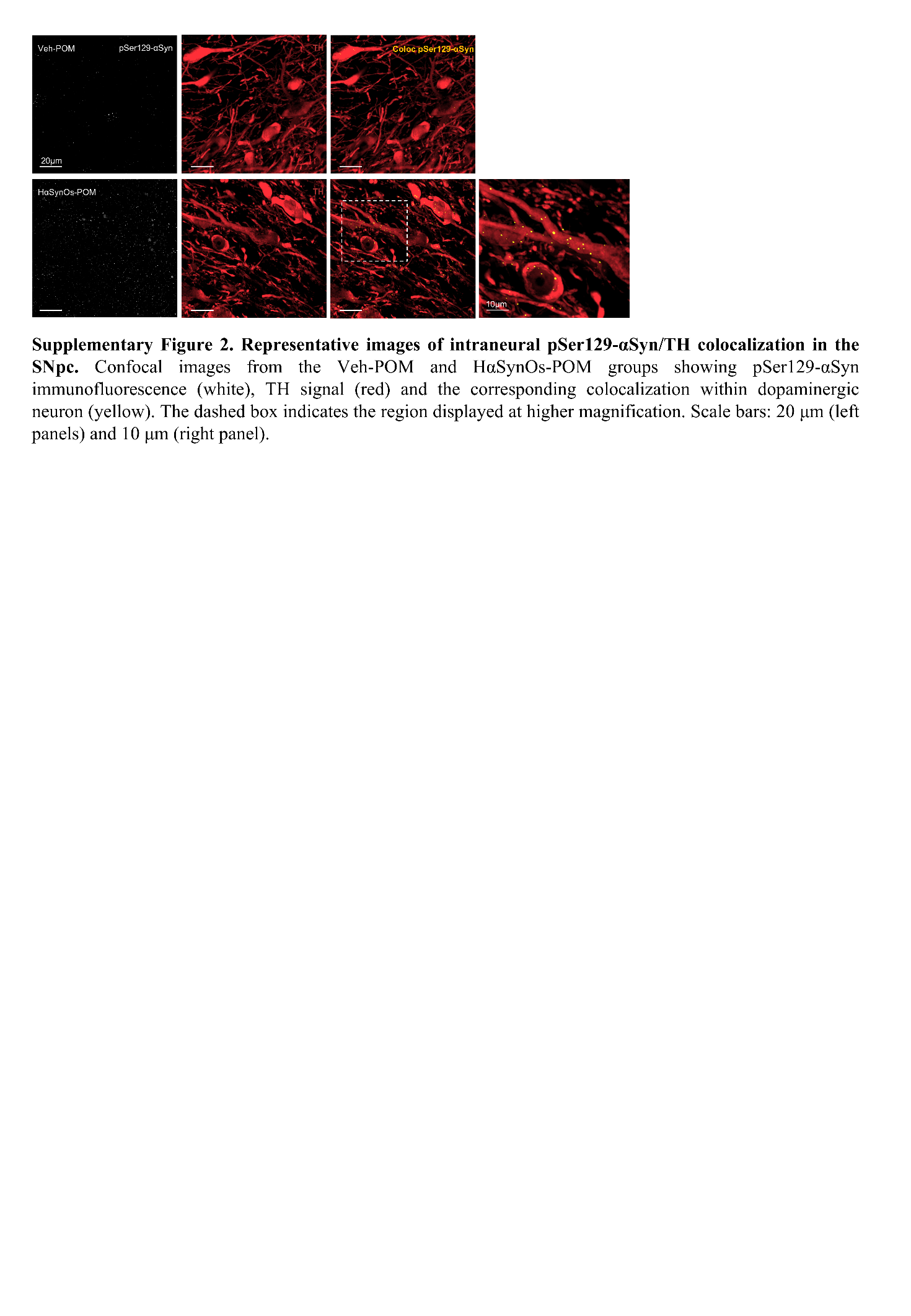

**Supplementary Figure 2. Representative images of intraneural pSer129-αSyn/TH colocalization in the SNpc.** Confocal images from the Veh-POM and HαSynOs-POM groups showing pSer129-αSyn immunofluorescence (white), TH signal (red) and the corresponding colocalization within dopaminergic neuron (yellow). The dashed box indicates the region displayed at higher magnification. Scale bars: 20 μm (left panels) and 10 μm (right panel).

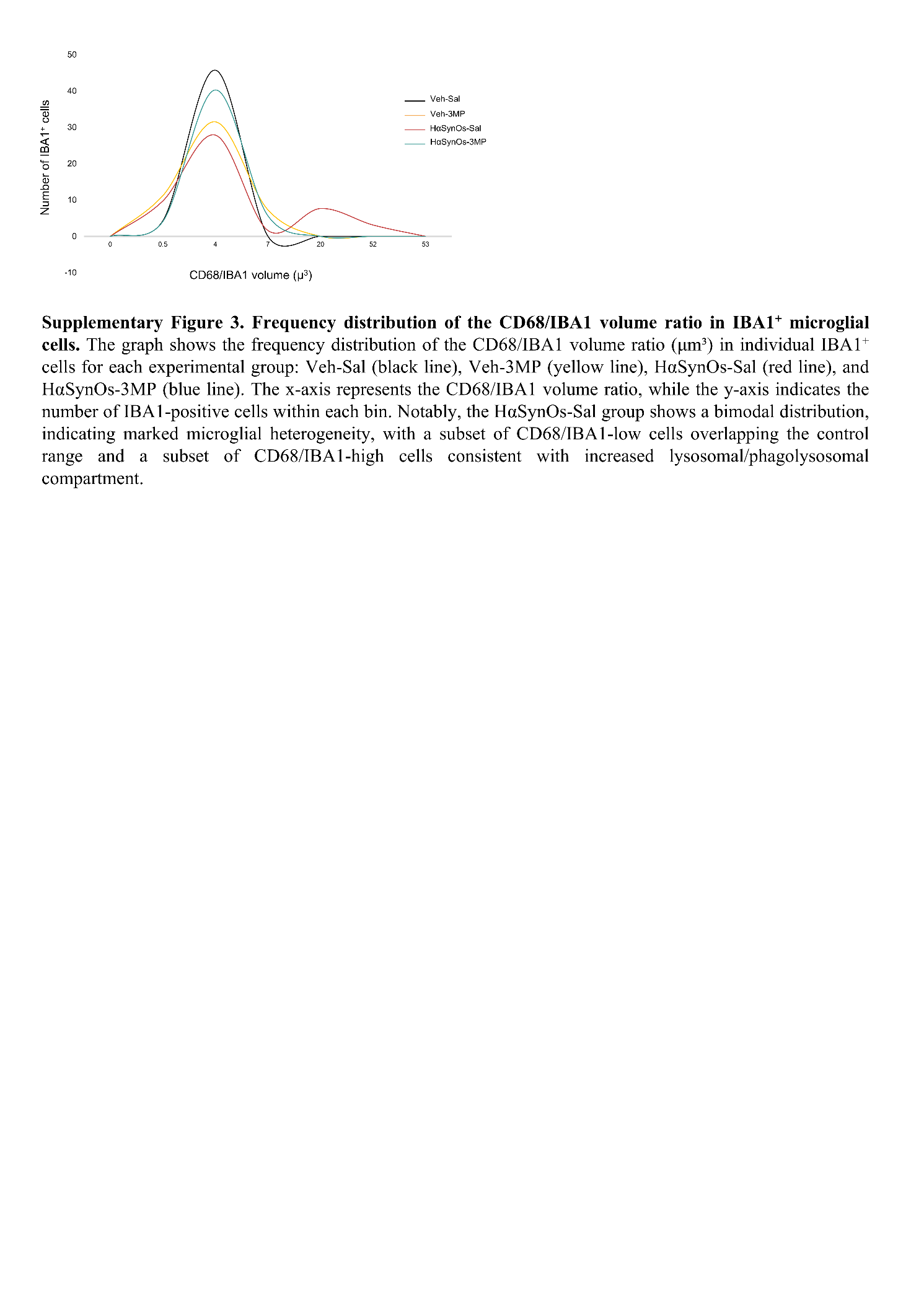

**Supplementary Figure 3. Frequency distribution of the CD68/IBA1 volume ratio in IBA1+ microglial cells.** The graph shows the frequency distribution of the CD68/IBA1 volume ratio (μm³) in individual IBA1+ cells for each experimental group: Veh-Sal (black line), Veh-3MP (yellow line), HαSynOs-Sal (red line), and HαSynOs-3MP (blue line). The x-axis represents the CD68/IBA1 volume ratio, while the y-axis indicates the number of IBA1-positive cells within each bin. Notably, the HαSynOs-Sal group shows a bimodal distribution, indicating marked microglial heterogeneity, with a subset of CD68/IBA1-low cells overlapping the control range and a subset of CD68/IBA1-high cells consistent with increased lysosomal/phagolysosomal compartment.
